# A bioenergetic basis for multiorgan dysfunction in sepsis

**DOI:** 10.1101/2025.06.12.659280

**Authors:** Elisa Jentho, Temitope Wilson Ademolue, Hessel Peters-Sengers, Joe M. Butler, Lin- Lin Xu, Sonia Trikha Rastogi, Jamil Kitoko, Sara Pagnotta, Bernhard Drotleff, Pedro Faísca, Valentyna Polishchuk, Miguel Mesquita, Paul Horn, Uta Barbara Metzing, Sílvia Cardoso, Gilles Mithieux, Michael Bauer, Osoul Chouchane, Sebastian Weis, Gianni Panagiotou, Christian von Loeffelholz, Tom van der Poll, Miguel P. Soares

**Affiliations:** Gulbenkian Institute for Molecular Medicine, Rua da Quinta Grande 6, 2780-156 Oeiras, Portugal; Jena University Hospital, Department of Anesthesiology and Intensive Care Medicine, Friedrich-Schiller-University Jena, Am Klinikum 1, 07747 Jena, Germany; Center for Infection and Molecular Medicine, Amsterdam University Medical Center, Academic Medical Center, University of Amsterdam, Meibergdreef 9, 1105 AZ, Amsterdam, The Netherlands; The Amsterdam Institute for Infection and Immunity, Amsterdam University Medical Center, Meibergdreef 9, 1105 AZ, Amsterdam, The Netherlands; Department of Epidemiology and Data Science, Amsterdam University Medical Center, Vrije Universiteit Amsterdam, Amsterdam, The Netherlands; Department of Microbiome Dynamics, Leibniz Institute for Natural Product Research and Infection Biology - Hans Knöll Institute, Jena, Germany; Metabolomics Core Facility, European Molecular Biology Laboratory, Meyerhofstraße 1, 69117 Heidelberg, Germany; Charité – Universitätsmedizin Berlin - Department of Hepatology and Gastroenterology, Campus Virchow-Klinikum and Campus Charité Mitte, Charitéplatz 1, 10117 Berlin, Germany; Charité – Universitätsmedizin Berlin – Berlin Institute of Health at Charité, BIH Biomedical Innovation Academy, Kapelle-Ufer 2, 10117 Berlin, Germany; Jena University Hospital, Department of Trauma, Hand and Reconstructive Surgery, Friedrich-Schiller-University Jena, Am Klinikum 1, 07747 Jena, Germany; Université Claude Bernard Lyon, INSERM, 43 Boulevard du 11 Novembre 1918, 69622 Villeurbanne cedex, Lyon, France; Center for Sepsis Control and Care, Friedrich-Schiller-University Jena, Am Klinikum 1, 07747 Jena, Germany; Jena University Hospital, Institute for Infectious Disease and Infection Control, Friedrich-Schiller-University Jena, Am Klinikum 1, 07747 Jena, Germany; Leibniz Institute for Natural Product Research and Infection Biology - Hans Knöll Institute, Jena, Germany; Faculty of Biological Sciences, Friedrich Schiller University, Jena, Germany; Department of Medicine, The University of Hong Kong, Hong Kong S.A.R., China; Division of Infectious Diseases, Amsterdam University Medical Center, University of Amsterdam, Meibergdreef 9, 1105 AZ, Amsterdam, The Netherlands

## Abstract

Sepsis is a life-threatening multiorgan dysfunction that develops from a maladaptive host response to infection^1^. With an estimated 49 million cases per year and ∼11 million related deaths^2^, sepsis is a global WHO health priority^3^. Failure to overcome sepsis morbidity and lethality^4,5^ calls for alternative therapeutic approaches^6–8^. Here we report that adipocyte lipolysis is vital to prevent the pathogenesis of sepsis in mice. This protective response is evolutionary conserved, producing a plasma lipidomic profile^9,10^ that reflects on the severity of clinical sepsis. Mechanistically, adipocyte lipolysis fuels energy metabolism to sustain adaptive thermoregulation to infection, via insulin production and insulin receptor (INSR) signaling in adipocytes. This metabolic-based defense strategy does not impact on bacterial burden, establishing disease tolerance to infection^11–14^. In conclusion, adipocyte lipolysis induces insulin to rewire energy metabolism and support organ function in response to infection.

---

The pathogenesis of sepsis^15^ is associated with an unfettered inflammatory response^16^ that induces various sickness behaviors^17,18^, including illness-induced anorexia, which reduces food (*i.e.,* nutrient) intake^19^. The infected host develops a starvation-like response^20^, characterized initially by a transient state of hyperglycemia fueled by the induction of hepatic glucose production (HGP)^21^. This is followed by a sustained reduction of glycemia eventually leading to hypoglycemia due to repression of HGP^21–25^. Consistent with this starvation-like response, most sepsis patients develop hyperglycemia due to increased HGP and peripheral insulin resistance, with a smaller fraction developing hypoglycemia ^26^.

Rewiring of host glucose metabolism is protective against bacterial^21,22^ and protozoan^23,27^ infection in mice. This metabolic-based defense strategy acts irrespectively of the host’s pathogen burden, establishing disease tolerance to infection^11,13,14^.

Insulin, the master regulator of glucose metabolism, acts via the INSR^28^ to reduce glycemia, via different mechanisms that include the repression of HGP^29^ and the induction of cellular glucose uptake^28,30^. The development of life-threatening hypoglycemia is prevented by a controlled repression of INSR signaling, a phenomenon known as insulin resistance^31^. To what extent insulin regulates disease tolerance to infection is not clear^23^. Of note, insulin is used as a standard therapeutic intervention in hyperglycemic control of sepsis patients^32,33^.

The starvation-like response to infection is associated with the induction of adipocyte lipolysis^17,18^, as illustrated in mice^34–37^ and humans^38^, including in critically ill patients^39,40^. Here we addressed whether adipocyte lipolysis and its regulation by insulin^41,42^ represents an adaptive or maladaptive response to sepsis.

C57BL/6J mice developed prototypical sickness behaviors in response to polymicrobial infection, induced by cecal ligation puncture (CLP; ∼50% lethality; LD_50_) (*Extended Data Fig. 1A*). These included reduction of physical activity, responsiveness to stimulus and consciousness (*Extended Data Fig. 1B*) as well as illness-induced anorexia (*Extended Data Fig. 1C*). This was associated with a transient reduction of body weight (*i.e.,* wasting), glycemia and core body temperature (*i.e.,* hypothermia)(*Extended Data Fig. 1C*), regaining physiologic levels within 7-8 days post-infection, with the exception of body weight (*Extended Data Fig. 1C*)

To determine the contribution of illness-induced anorexia to the development of wasting, hypoglycemia and hypothermia, we pair-fed^43^ non-infected (sham operated) and infected (CLP; LD_50_) mice to normalize their food intake. Non-infected mice developed a wasting response when pair-fed to infected mice (*Extended Data Fig. 1C*). This suggests that illness-induced anorexia is the main driving force in the wasting response of infected mice.

In contrast, pair-fed control mice did neither develop hypoglycemia nor hypothermia (*Extended Data Fig. 1C*), suggesting that the illness-induced anorexia is not the main driving force for the development of hypoglycemia and hypothermia in response to infection. We infer that infection elicits a host metabolic response mechanistically distinct from the starvation response^44^.

C57BL/6J mice subjected to CLP (LD_50_) presented a progressive reduction of epididymal (e) white adipose tissue WAT (eWAT) (*Extended Data Fig. 2A, B*), subcutaneous WAT (scWAT) (*Extended Data Fig. 2C, D*) and brown adipose tissue (BAT) (*Extended Data Fig. 2E, F*) mass. This is consistent with adipocyte lipolysis driving the wasting response to infection^8^.

The wasting response of infected mice was propelled by a reduction of lipid vacuole size (*Extended Data Fig. 2B, D, F*). This reduction was associated with the induction of lipolysis^45–48^, as reveled by the induction of adipocyte triglyceride lipase (ATGL) expression and the activation of the hormone sensitive lipase (HSL) (*i.e.,* Ser660 phosphorylation) in eWAT (*Fig. 1A,B*; *Extended Data Fig. 3A*, *Video 1&2*) and BAT (*Extended Data Fig. 3B,C*, *Video 3&4*).

**Figure 1:**
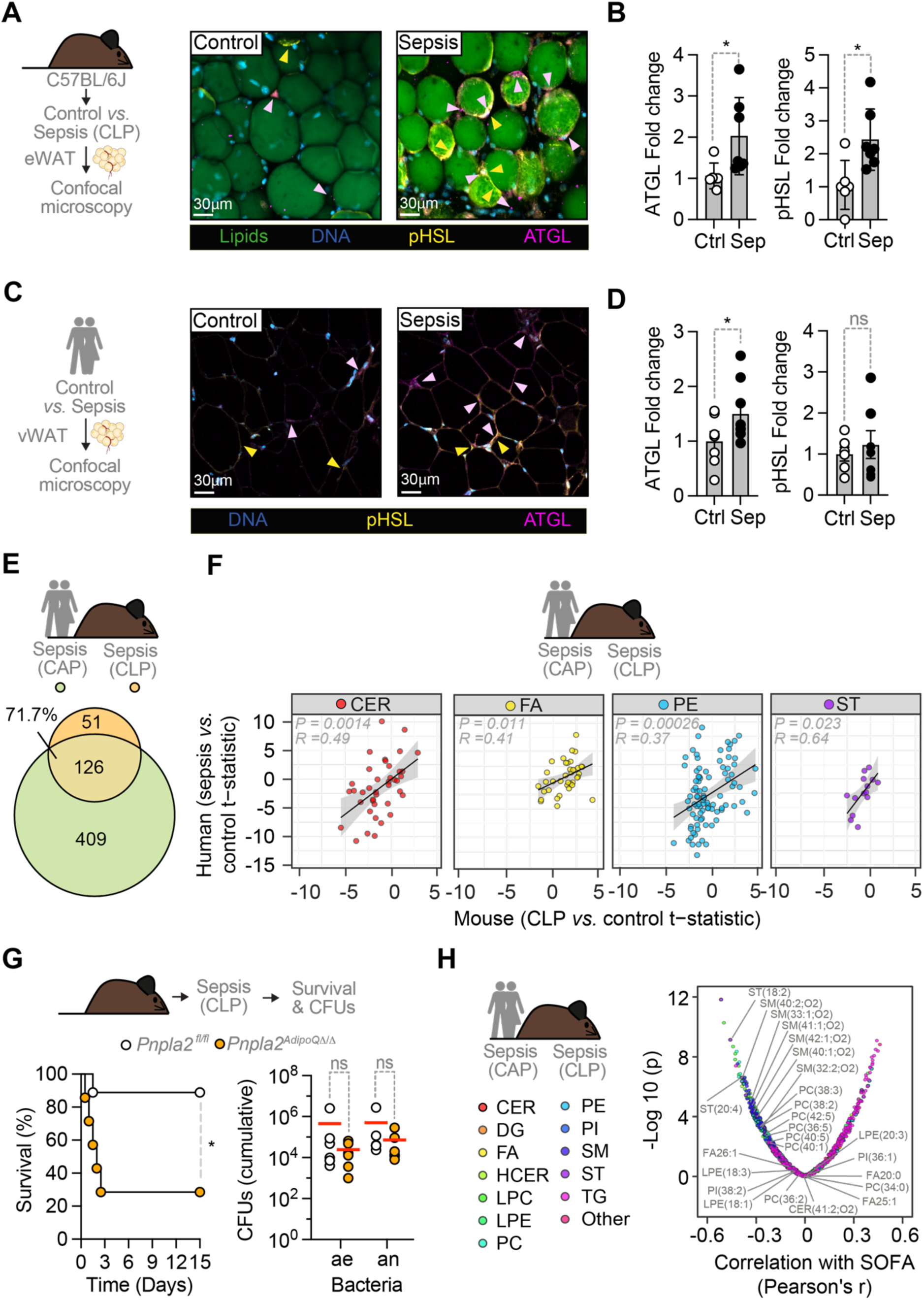
Adipocyte lipolysis is an adaptive response to infection in mice and humans. **A)** Immunofluorescence imaging of neutral lipids (BODIPY; green), DNA (DAPI; blue), adipose triglyceride lipase (ATGL; anti-ATGL antibody with Alexa Fluor® 647; magenta) and phospho- hormone sensitive lipase (pHSL; anti-pHSL Ser660 antibody with Alexa Fluor® 568; yellow) of eWAT from C57BL/6J mice, 24hrs after sham operation (Control, Ctrl) *vs.* CLP (Sepsis). **B)** Quantification of ATGL and p-Ser660-HSL. Data pooled from N=5-7 *per* group, represented as mean ± SEM, from 23 different areas in the same images, from 23 images from 2 independent experiments with similar trend. Circles represent individual mice. P-values calculated by Student’s t-test, *p≤0.05. **C)** Immunofluorescence imaging as in (A) without neutral lipids, from visceral WAT of control vs. septic patients. **D)** Quantification of ATGL (N=7- 8 individuals) and p-Ser660-HSL (N=6-7 individuals). Data represented as mean ± SEM, from 23 different areas in the same images, pooled from 21 image(s). Circles represent different individuals. P-values calculated by one-sided Student’s t-test, *p≤0.05. **E)** Venn-diagram of overleaping lipid metabolites significantly regulated in plasma from septic patients with community acquired pneumonia (CAP) (green, N=169) and septic mice (orange, N=3) *vs.* human (N=48) controls and mouse (N=4), respectively. **F)** Correlation of t-statistics for ceramides (CER), fatty acids (FA), phosphatidylethanolamine (PE) and sterols (ST) in plasma from CAP septic (N=169) vs. controls (N=48) patients and septic (CLP; N=3) vs. control (N=4) mice. **G)** Survival of septic (CLP) *Pnpla2^fl/fl^* (N=9) *vs. Pnpla2^AdipoQ11/11^* (N=7) mice. Data pooled from 3 independent experiments, with similar trend. P-values calculated by Log-rank (Mantel- Cox) test, *p<0.05. Cumulative colony forming units (CFU; ae: aerobe, an: anaerobe) from different organs *(Fig. S8B*), 24hrs after CLP (N=4-6 per genotype). Data pooled from 3 independent experiments, with similar trend. Circles represent individual mice. P-values calculated by Students t-test, ns – not significant, *p<0.05. **H)**. Correlation between SOFA severity score and lipid level in CAP septic patients; lipids were labelled when significantly regulated in both patients and mice, compared to respective control groups (Human cut off: BH adjusted < 0.05; Mouse cut off: top 25 downregulated). P-values were calculated by univariate linear regression.

We questioned whether the lipolytic response to infection is evolutionary conserved in mice and humans. In support of this hypothesis, peritoneal sepsis patients (*Table 1*)^49^ also presented an increase in adipocyte ATGL expression, but not HSL activation, in visceral WAT (vWAT), compared to non-septic controls undergoing open abdominal surgery (*Fig. 1C; Extended Data Fig. 3D*).

**Table 1:** Clinical characteristics of the control and peritoneal sepsis patients.

|  | **Control (n=10)** | **Sepsis (n=10)** | **p-value** |
| --- | --- | --- | --- |
| Age (years) | 58 [52-64] | 69.5 [61.5-74.2] | 0.037 |
| Sex (male) | 3/10 | 5/10 | 0.65 |
| BMI (kg/m<sup>2</sup>) | 21.8 [21.4-24.2] | 28.9 [25.6-31.6] | 0.023 |
| HbA<sub>1c</sub> (%) | 5.5 [5.3-5.5] | 5.8 [5.6-6.1] | 0.118 |
| Fasting Glucose (mmol/L) | 5.2 [4.9-5.3] | 5.2 [4.7-6.8] | 0.733 |
| Triacylglycerol (mmol/L) | 1.19 [0.89-1.54] | 2.30 [1.51-2.73] | 0.104 |
| Free Fatty Acids (mmol/L) | 0.67 [0.35-0.92] | 0.94 [0.55-1.04] | 0.290 |
| Total cholesterol (mmol/L) | 5.1 [4.6-5.4] | 2.0 [1.4-2.3] | <0.001 |
| LDL cholesterol (mmol/L) | 3.7 [2.92-3.97] | 0.5 [0.42-0.63] | <0.001 |
| HDL cholesterol (mmol/L) | 1.55 [1.2-1.77] | 0.17 [0.13-0.40] | 0.0028 |
| Creatinine (mmol/L) | 64.5 [59.5-71.2] | 249 [92-292] | 0.0011 |
| Bilirubin (mmol/L) | 8.5 [6.5-10.8] | 23 [15.2-29] | 0.0057 |
| CRP (mg/dL) | 2.25 [2-3.48] | 250 [181-287] | <0.001 |
| WBC (10<sup>9</sup>/L) | 5.65 [4.53-6.33] | 16.1 [12.1-19.9] | <0.001 |
| Platelet count (10<sup>9</sup>/L) | 222 [201-262] | 328 [200-445] | 0.345 |
| SOFA score | - | 8 [6-11] |  |
| APACHE II score | - | 20.5 [16.5-24.8] |  |

All parameters, except for sex, are given as median [IQR]. Statistical comparisons were performed using unpaired two-sided Wilcoxon-Test or $\chi^2$ -test.

C57BL/6J mice subjected to CLP (LD_50_) presented a transient increase in the concentration of free fatty acids (FA) and glycerol in plasma, with a tendency, *albeit* not significant for a reduction of triglycerides (TG), compared to non-infected controls (*Extended Data Fig. 4A*). High-resolution mass spectrometry/liquid chromatography^50^ confirmed that the plasma lipidomic profile of C57BL/6J subjected to CLP (LD_50_) was distinct from that of controls subjected to “sham” surgery or naïve controls, as assessed by principal component analysis (PCA) (*Extended Data Fig. 4B*).

Infected mice accumulated FA (*Extended Data Fig. 4C,D*) while lowering triglycerides (TG), ceramides (CER), hexoceramides (HCER), lysophosphatidylcholines (LPC), phosphatidylcholines (PC), phosphatidylethanolamines (PE) and sterols (ST) in plasma (*Extended Data Fig. 4C,D*), as compared to naïve controls. Other lipid metabolites such as diacylglycerides (DG), lysophosphatidylethanolamines (LPE), Phosphatidylinositols (PI), sphingomyelins (SM) and phosphatidylinositols (PI) were not changed in response to infection (*Extended Data Fig. 4C,D*), consistent with previously described^51^.

The notion that adipocyte lipolysis is an evolutionary conserved response to infection is further supported by the major overlap (71%; 126 lipid metabolites) of the significantly regulated lipid metabolites in plasma lipidomic profile of septic patients from a community acquired pneumonia cohort^10^ and septic mice (*Fig. 1E,F, Extended Data Fig. 5*). T-statistics for all individual lipid metabolites from septic humans and mice (*i.e.,* relative to controls) showed a significant positive correlation (*Extended Data Fig. 6A*), attributed to FA, CER, PE and ST (*Fig 1F*). Other lipid metabolites showed no or negative correlations (*Extended Data Fig. 6B*).

We tested the contribution of adipocyte lipolysis to the plasma lipidomic profile of septic mice in *Pnpla^2AdipoQ11/11^* mice, carrying a genetic loss-of-function of the patatin-like phospholipase domain-containing protein 2 gene (*Pnpla2;* encoding ATGL) specifically in adipocytes^52^. *Pnpla2^AdipoQ11/11^* mice subjected to CLP (LD_20_) presented a plasma lipidomic profile that was distinct from control *Pnpla2^fl/fl^*, as revealed by PCA (*Extended Data Fig. 7A*). This was accounted for by a reduction of circulating FA, CER, DG, HCER, LPE, PC, PE, SM, ST and TG in *Pnpla2^AdipoQ11/11^ vs.* control *Pnpla2^fl/fl^* mice (*Extended Data Fig. 7B,C*), suggesting that adipocyte lipolysis regulates the release and/or synthesis of these lipid metabolites in response to infection.

We followed up by asking whether the plasma lipidomic profile of *Pnpla2^AdipoQ11/11^* mice is associated with sepsis severity. *Pnpla2^AdipoQ11/11^* succumbed readily to polymicrobial sepsis, when subjected to the CLP LD_20_ in control *Pnpla2^fl/fl^* mice (*Fig. 1G*). This was also observed using contamination and infection (PCI) (*Extended Data Fig. 8A*), an experimental model of polymicrobial sepsis where mice are infected by the same inoculum.

The bacterial burden was indistinguishable between *Pnpla2^AdipoQ11/11^* mice subjected to CLP and that of *Pnpla2^fl/fl^*mice that survived the infection (*Fig. 1G*), as assessed for aerobic and anaerobic bacteria in different organs (*Extended Data Fig. 8B*). This suggests that adipocyte lipolysis is essential to establish disease tolerance to polymicrobial infection.

We then addressed whether the plasma lipidomic profile of lethal sepsis in *Pnpla2^AdipoQ11/11^* mice relates to sepsis severity in community acquired pneumonia (CAP) patients^10^. Consistent with this hypothesis, most lipid metabolites downregulated in septic *Pnpla2^AdipoQ11/11^ vs.* control *Pnpla2^fl/fl^* mice overlapped with those downregulated in septic *vs.* control patients (79.3%; 180 out of 227 down-regulated lipid metabolites) (*Extended Data Fig. 9A,B*). Of note, one lipid metabolite (PC34:1) was upregulated in both infected *Pnpla2^AdipoQ11/11^ vs. Pnpla2^fl/fl^* mice and in CAP sepsis *vs.* control patients (*Extended Data Fig. 9A,B*), consistent with previous findings^51^. Among the top 25 lipid metabolites most significant downregulated in infected *Pnpla2^AdipoQ11/11^ vs. Pnpla2^fl/fl^* mice, 56% (14 PC, SM and ST) were negatively correlated with clinical sepsis severity, as assessed by Sequential Organ Failure Assessment (SOFA) score in community acquired pneumonia patients (*Fig 1H, Extended Data Fig.10*). This is consistent with adipocyte lipolysis generating a plasma lipidomic profile that predicts sepsis severity in mice and humans.

To explore whether adipocyte lipolysis is required to overcome the energy deficit imposed by illness induced anorexia, *Pnpla2^AdipoQ11/11^* mice were pair-fed to infected *Pnpla2^fl/fl^* mice. Pair-feeding was not lethal to *Pnpla2^AdipoQ11/11^* mice (*Extended Data Fig. 11*), suggesting that illness induced anorexia does not explain *per se* the lethal outcome of infection in *Pnpla2^AdipoQ11/11^* mice.

We proceeded to ask whether the protective effect of adipocyte lipolysis is mediated by the immunoregulatory effects attributed to lipolytic products^53^. In keeping with this notion, the levels of circulating interleukin (IL)-1β, previously implicated in the pathogenesis of sepsis^16^ was higher in infected *Pnpla2^AdipoQ11/11^ vs.* control *Pnpla2^fl/fl^* mice (*Extended Data Fig. 12*). This was also observed for other cytokines, including IL-1α, IL-10, IL-12, IL-23 and C-C Motif Chemokine Ligand 2 (CCL2/MCP-1) in infected *Pnpla2^AdipoQ11/11^ vs. Pnpla2^fl/fl^* mice (*Extended Data Fig. 12*). This increase in cytokine production was however modest, and therefore not likely to account, *per se*, to the protective effect of adipocyte lipolysis against sepsis.

The lethal disease course of infected *Pnpla2^AdipoQ11/11^* mice was associated with the development of multiorgan dysfunction, illustrated by higher accumulation of circulating lactate dehydrogenase (LDH), aspartate aminotransferase (AST; liver damage), alanine aminotransferase (ALT; liver damage), urea (kidney dysfunction) and creatinine phosphokinase (CPK; muscle and heart damage)(*Extended Data Fig. 13A*), compared to infected control *Pnpla2^fl/fl^*mice. Multiorgan injury was not reflected histologically (*Extended Data Fig. 12B*), consistent with clinical sepsis^6–8^.

The cardiac and hepatic metabolomic profile of infected *Pnpla2^AdipoQ11/11^* mice was distinct from that of infected-*Pnpla2^fl/fl^* control mice, as determined by PCA analyzes of untargeted metabolomics (*Fig. 2A*). This was not the case however, for the brain (*Fig. 2A*).

**Figure 2:**
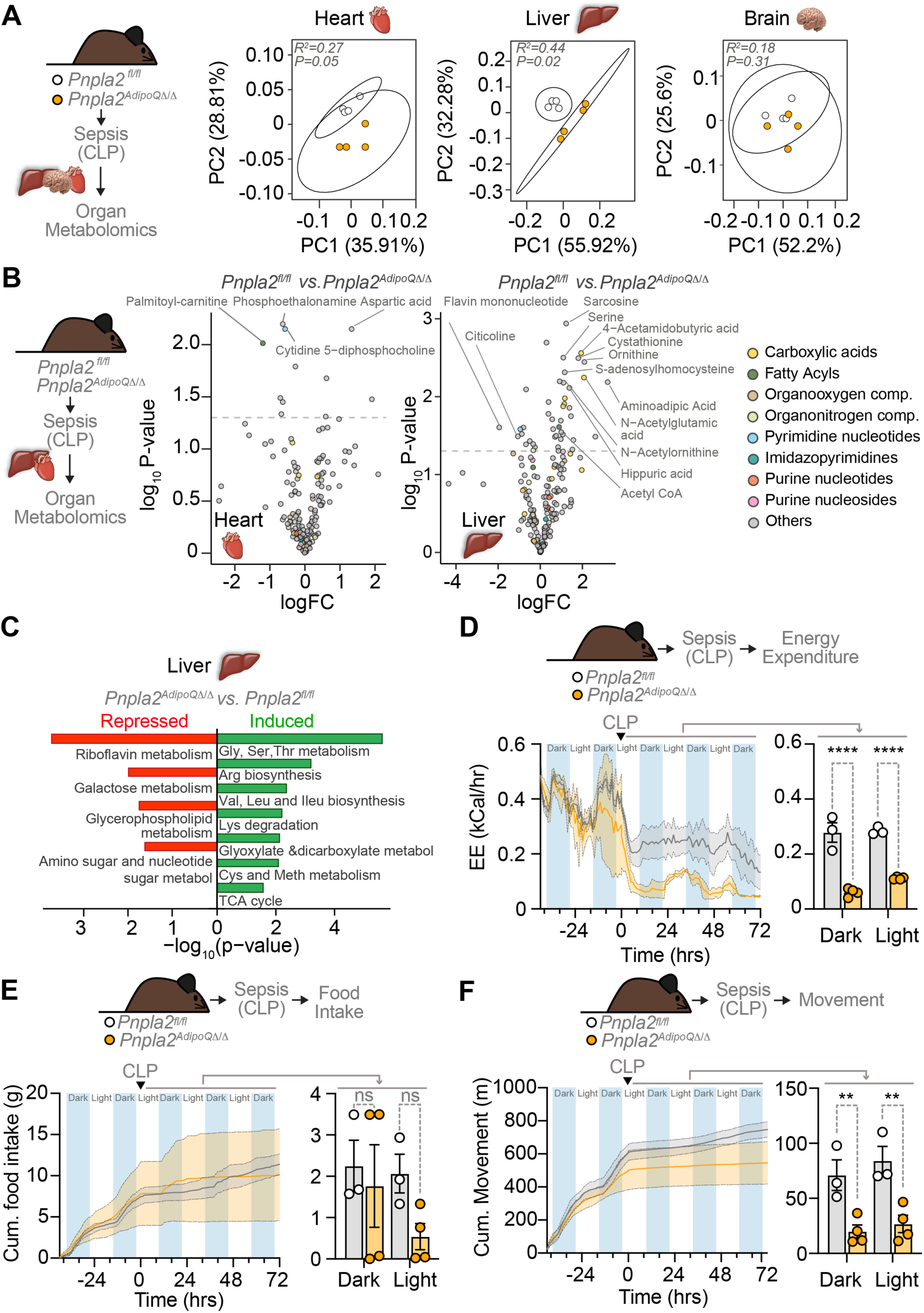
Adipocyte lipolysis supports organ function and energy metabolism in response to infection. **A)** PCA of untargeted metabolomics from heart, brain and liver of *Pnpla2^AdipoQ11/11^* (N=4) *vs. Pnpla2^fl/fl^* (N=4) mice, 24hrs after CLP (Sepsis). **B)** Differential abundance analysis of metabolites in heart and liver from same mice in (A). Significant metabolites considered as those with a P<0.05 and absolute fold change >1. **C)** Significantly regulated metabolic pathways, defined by MetaboAnalyst in the liver of the same mice from (A,B). **D)** Energy expenditure (EE) of *Pnpla2^AdipoQ11/11^* (N=4) *vs. Pnpla2^fl/fl^* (N=3) mice, at steady state or after CLP (dotted line; 0 hrs). Right panel shows the mean EE of daily averages at dark and light cycles after CLP, represented as mean ± SEM. Circles represent individual mice. **(E)** Cumulative food intake of same mice as (D). Right panel shows the mean of daily food intake averages at dark and light cycle after CLP, as mean ± SEM. Circles represent individual mice. **(F)** Cumulative movement of same mice as (D). Right panel shows the mean movement of daily averages at dark and light cycle after CLP, as mean ± SEM. Circles represent individual mice. P-values in (D,E,F) calculated by Two-Way ANOVA with Šídák’s multiple comparisons test, *p≤0.05, ****p≤0.0001; ns – not significant. Data from one experiment.

Next, we inquired whether adipocyte lipolysis rewires the metabolic function of vital organs. Consistent with this hypothesis we found that the hepatic metabolomic profile of infected *Pnpla2^AdipoQ11/11^* mice showed profound differences in lipid metabolism (*i.e.,* serine, acetyl-CoA, citicoline), redox balance (*i.e.,* flavin mononucleotide; FMN), autophagy (*i.e.,* sarcosine), xenobiotic detoxification (*i.e.,* hippuric acid, FMN), nitrogen metabolism [*i.e.,* ornithine, N- acetylglutamate (NAG), N-acetylornithine] and methylation [*i.e.,* sarcosine, S-Adenosyl-L- homocysteine (SAH)], compared to control infected-*Pnpla2^fl/fl^* mice (*Fig. 2B,C*). This suggests that adipocyte lipolysis is essential to rewire hepatic metabolic function in response to infection.

Adipocyte lipolysis supports energy metabolism in response to energy shortage^54^ entertaining the hypothesis that this is also the case during severe infection. In support of this hypothesis, *Pnpla2^AdipoQ^*^11/11^ mice developed a life-threating reduction of energy expenditure (EE; total amount of calories used)^55^ in response to infected, compared to the viable reduction EE in control infected *Pnpla2^fl/fl^* mice (*Fig. 2D*). This EE collapse occurred under the same food intake (*Fig. 2E*) and motor activity (*Fig. 2F*), suggesting that adipocyte lipolysis is essential to sustain the lower threshold of EE required to survive infection. Of note infected *Pnpla2^AdipoQ^*^11/11^ mice presented a proportional reduction of oxygen consumption (VO_2_) (*Extended Data Fig. 14A*) and CO_2_ production (VCO_2_) (*Extended Data Fig. 14B*), not reflected by a concomitant reduction of respiratory quotient (Rq; VO_2/_VCO_2_) (*Extended Data Fig. 14C*).

The lethal outcome of infection in *Pnpla2^AdipoQ^*^11/11^ mice was associated with a life-threatening reduction of BAT thermogenesis (*Fig. 3A,B*) and body temperature (*Fig. 3A-C*), as well as with higher heat dissipation (*Fig. 3B*), compared to control infected *Pnpla2^fl/fl^* mice (*Fig. 3A-C*). This suggests that adipocyte lipolysis “guards” the lower threshold of the adaptive reduction of BAT thermogenesis in response to bacterial infection^25^. Moreover, this is also consistent with adipocyte lipolysis^56^ fueling BAT thermogenesis^56,57^.

**Figure 3:**
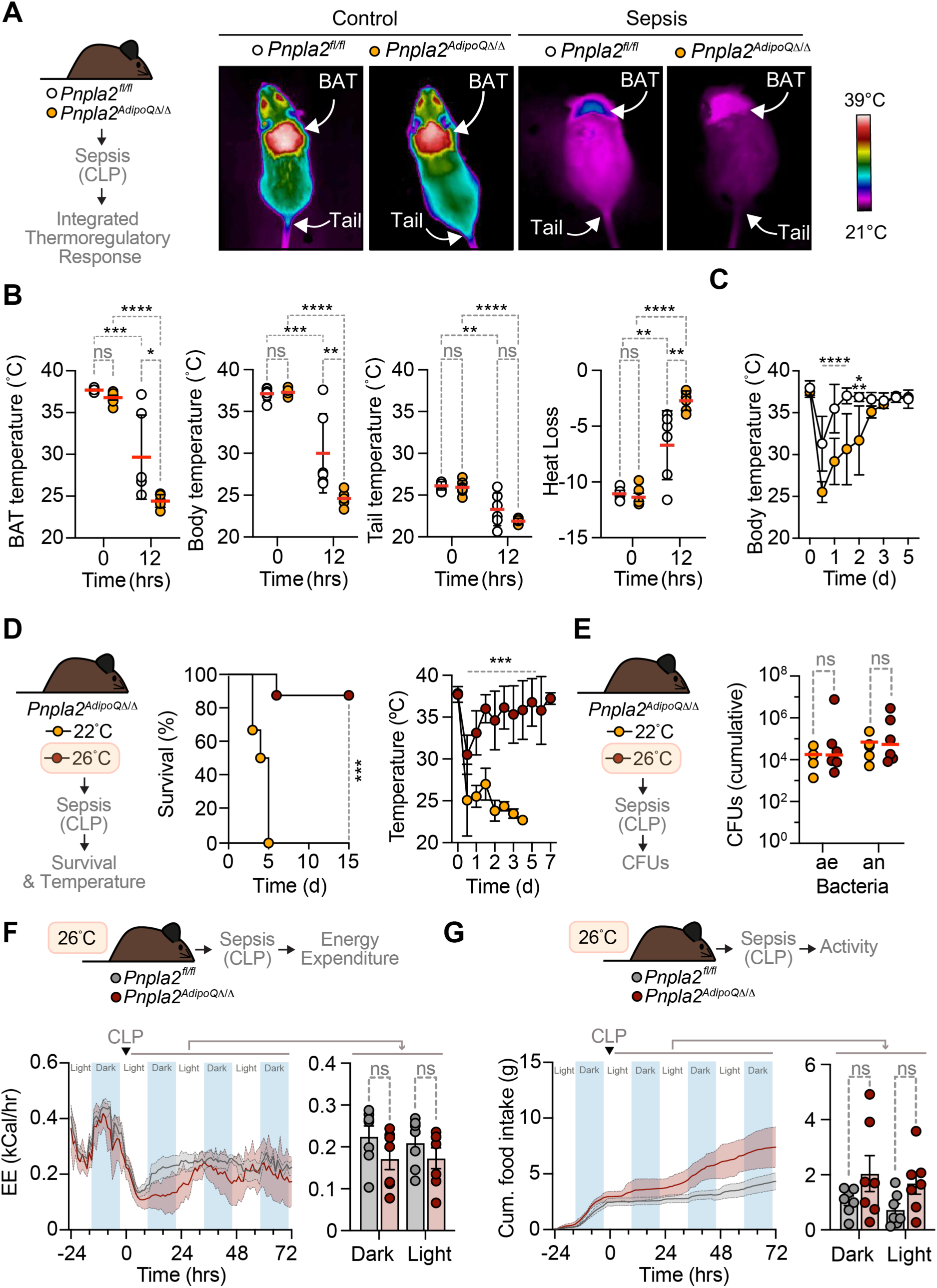
Adipocyte lipolysis supports the lower threshold of the adaptive hypothermic response to infection. **A)** Representative infrared images of *Pnpla2^AdipoQ11/11^* and *Pnpla2^fl/fl^* mice at steady state (Controls) *vs.* 12hrs after CLP (Sepsis). **B)** Quantification of brown adipose tissue (BAT) temperature, core body temperature, tail temperature and heat loss (ι1 of tail base and core body temperature). Data represented as mean (red bars) ± SEM (black whiskers), from 23 different images from the same mouse (N=6 mice *per* genotype), pooled from 2 independent experiments, with similar trend. P-values calculated by Two-Way ANOVA with Šídák’s multiple comparisons test. Circles represent individual mice. **C)** Core body temperature over time of *Pnpla2^AdipoQ11/11^* (N=7) *vs. Pnpla2^fl/fl^*(N=9) mice subjected to CLP (same mice as Fig. 1G). Data represented as mean ± SD, pooled from 3 independent experiments, with similar trend. P-values calculated by Mixed-Effect-Model analysis with Šídák’s multiple comparisons test. **D)** Survival of *Pnpla2^AdipoQ11/11^* mice subjected to CLP and maintained at 22-23°C (N=6) *vs.* 26-27°C (N=8). Data pooled from 2 independent experiments, with similar trend. P values calculated by Log-rank (Mantel-Cox) test. Right panel shows core body temperature from the same mice, represented as mean ± SD. P-values calculated by Two-Way ANOVA with Šídák’s multiple comparisons test. **E)** Cumulative colony forming units (CFU; ae: aerobe, an: anaerobe) from different organs (*Fig. S15E*) of *Pnpla2^AdipoQ11/11^* mice maintained at 22-23°C (N=6) *vs.* 26-27°C (N=8), 24h after CLP. Data represented as mean ± SD, pooled from 2 independent experiments (N=4-6 *per* experimental condition), with similar trend. P-values calculated by Students t-test. **F)** Energy expenditure (EE) of *Pnpla2^AdipoQ11/11^* (N=7) and *Pnpla2^fl/fl^* (N=7) mice, at steady state or after CLP (dotted line; 0 h), maintained at 26-27°C. Right panel shows EE of daily averages at dark and light cycle, after CLP, represented as mean ± SEM, pooled from 2 independent experiments, with similar trend. Circles correspond to individual mice. **(G)** Cumulative movement of same mice as (F). Right panel shows the mean movement of daily averages at dark and light cycle after CLP, represented as mean ± SEM. Circles correspond to individual mice. P-values in (F,G) calculated by Two-Way ANOVA with Šídák’s multiple comparisons test. ns – not significant., *p≤0.05; **p≤0.01; ***p≤0.001; ****p≤0.0001.

Mice allocate ∼1/3 of EE to support thermoregulation^25,58,59^ under standard (∼22°C) husbandry conditions, reducing heat dissipation and re-allocating EE towards other vital functions^25,58,59^ under thermoneutral (∼30°C) husbandry conditions. *Pnpla2^AdipoQ11/11^* mice survived polymicrobial infection when placed under husbandry conditions in the lower range of thermoneutrality (*i.e.,* ∼26°C), as compared to the lethal outcome of infection of *Pnpla2^AdipoQ11/11^* mice at standard conditions (*Fig. 3D*). This was associated with a control of hypothermia within a viable ∼30°C threshold, compared to the life-threatening ∼25°C threshold in *Pnpla2^AdipoQ11/11^* mice maintained at ∼22°C (*Fig. 3D*). Moreover, infected *Pnpla2^AdipoQ11/11^* mice maintained at ∼26°C restored EE (*Fig. 3F*), VO2 consumed (*Extended Data Fig. 15A*), VCO2 produced (*Extended Data Fig. 15B*) and Rq (*Extended Data Fig. 15C*) to the lower viable thresholds of infected *Pnpla2^fl/fl^* mice maintained at standard ∼22°C (*Fig. 3F,G*, *Extended Data Fig. 15B-D*). This regain of EE was not attributed to a putative increase in food intake (*Fig. 3E*) or a decrease in motor activity (*Extended Data Fig. 15D*). This suggests that adipocyte lipolysis is essential to “guard” the lower threshold of the adaptive hypothermic response to infection.

The bacterial burden of *Pnpla2^AdipoQ11/11^* mice that survive CLP when maintained at ∼26°C was indistinguishable from that is *Pnpla2^AdipoQ11/11^* mice that succumb to CLP when maintained at ∼22°C, as assessed for aerobic and anaerobic bacteria in different organs (*Fig. 3E, Extended Data Fig. 15E*). This shows that adipocyte lipolysis “guards” the lower threshold of the adaptive hypothermic response that establishes disease tolerance to infection^25^

Regulation of glucose metabolism plays a central role in disease tolerance to bacterial infection^21,22,60^, entertaining the hypothesis that disease tolerance relies on a crosstalk between glucose and lipid metabolism^61^. C57BL/6J mice increased glycemia within 2h of CLP (LD50), regaining basal levels within 6h (*Fig. 4A; Extended Data 16A*), consistent with described^21^. This was followed by a transient increase in insulinemia, peaking at 3h and regaining basal levels within 6h (*Fig. 4A; Extended Data 16B*). Activation (*i.e.,* phosphorylation) of protein kinase B (PKB/AKT) in BAT, 12-24h after CLP (*Fig. 4B*), suggests that the increase of insulinemia is associated with insulin receptor (INSR) signaling in response to infection.

**Figure 4:**
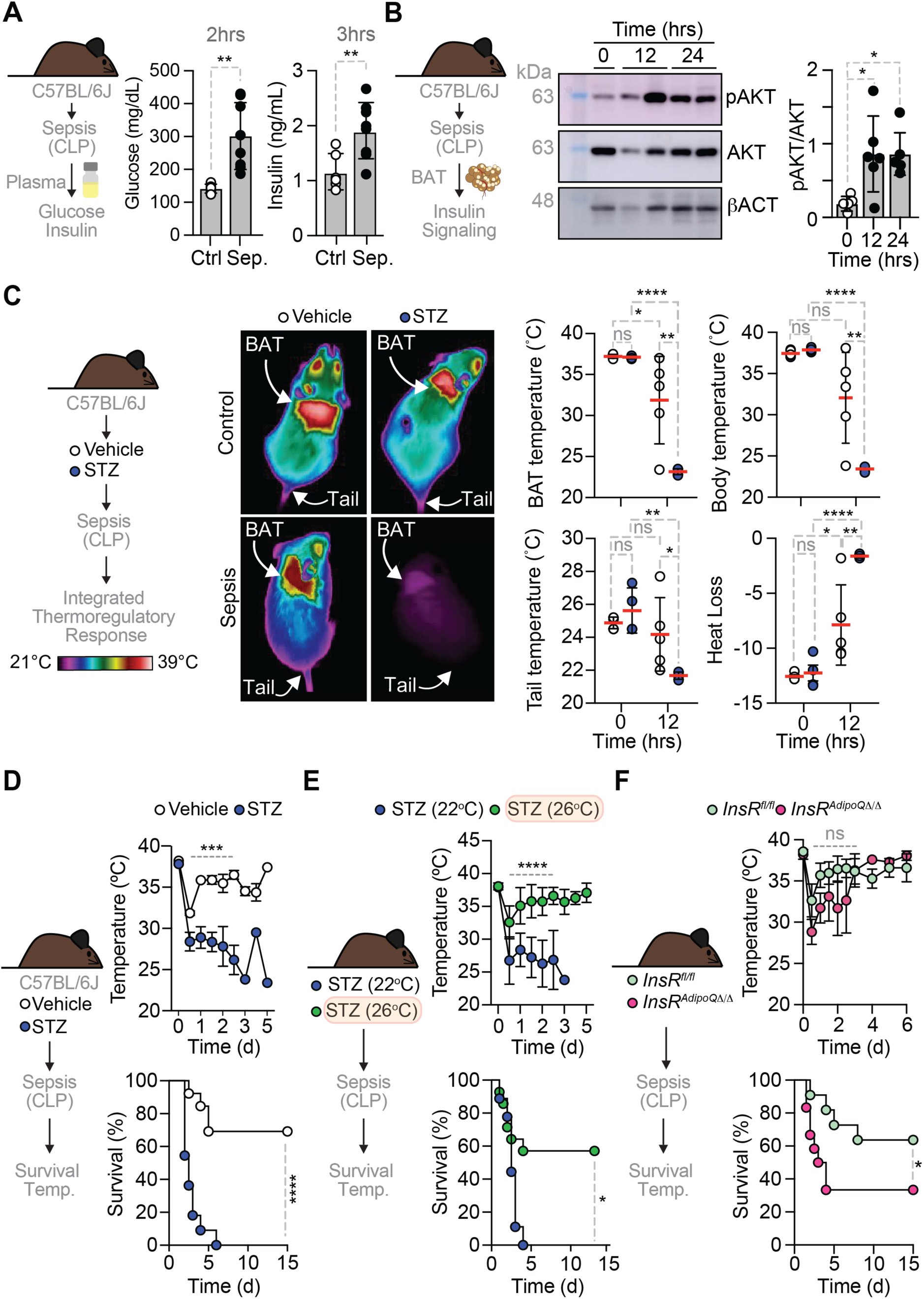
Insulin regulates the adaptive hypothermic response to infection. **A)** Glycemia and insulinemia of C57BL/6J mice 2 (glucose) and 3 (insulin) hours after a CLP (Sepsis; Sep.; N=8) or sham operation (Control; Ctr. N=6-8). Data represented as mean ± SD, from one experiment. P-values calculated by Students t-test. Circles represent individual mice. **B)** Representative western blot form BAT of C57BL/6J mice at steady state (0h), 12 and 24hrs after CLP (Sepsis). Right panel shows quantification of pAKT over AKT protein. Data represented as mean ± SD (N=6), from 2 independent experiments with same trend. P-values calculated by One-Way ANOVA with Dunnett multiple comparison test. **C)** Representative infrared images of C57BL/6J mice receiving PBS (vehicle; N=5) or Streptozocin (STZ; N=4) at steady state (Control) *vs.* 12hrs after CLP (Sepsis). Right panels show quantification of brown adipose tissue (BAT) temperature, core body temperature, tail temperature and heat loss (ι1 of tail base and core body temperature). Data represented as mean (red bars) ± SD (black whiskers) from ≥3 different images from the same mouse (N=4-5 mice *per* experimental condition), from one experiment. P-values calculated by Two-Way ANOVA with Šídák’s multiple comparisons test. Circles represent individual mice. **D)** Core body temperature and survival of C57BL/6J mice receiving PBS (vehicle; N=13) or Streptozocin (STZ; N=11) subjected to CLP (Sepsis). Data in upper panel represented as mean ± SD. **E)** Core body temperature and survival of C57BL/6J mice receiving Streptozocin (STZ) and subjected to CLP (Sepsis) at 22°C (N=9) or 26°C (N=14). Data in upper panel represented as mean ± SD. **F)** Core body temperature and survival of *InsR^AdipoQ11/11^* (N=12) *vs. InsR^fl/fl^* (N=11) mice subjected to CLP (Sepsis). Data in upper panel represented as mean ± SD. Data in (D-F) pooled from 3 independent experiments, with similar trend. P values in (D-F) calculated by Mixed-Effect-Model analysis with Šídák’s multiple comparisons test (upper panel) or Log-rank (Mantel-Cox) test (lower panel).

To test whether insulin regulates the adaptive hypothermic response that establishes disease tolerance to infection, insulin-producing pancreatic β-cells were ablated pharmacologically by Streptozocin (STZ)^23^. As expected^23^, C57BL/6J mice that received STZ became hyperglycemic (*Extended Data Fig. 16C*). This, however, did not impair the transient induction of hypoglycemia in response to CLP (*Extended Data Fig. 16C*), suggesting that the hypoglycemic response to infection is not driven by insulin.

C57BL/6J mice that received STZ failed to sustain BAT thermogenesis in response to infection (*Fig. 4C, Video 5,6*), developing life-threatening hypothermia, associated with increased heat dissipation (*Fig. 4C,D*). All C57BL/6J mice that received STZ succumbed to infection, compared to the ∼30% lethality (CLP; LD30) in controls (*Fig. 4D*). Mice that received STZ were protected from the lethal outcome of polymicrobial infection when maintained at ∼26°C (*Fig. 4E*). This was associated with a controlled reduction of body temperature to a viable ∼32°C threshold, compared to life-threatening ∼26°C threshold under standard ∼22°C husbandry conditions (*Fig. 4E*). It follows that insulin is essential to support BAT thermogenesis and “guard” the lower threshold of the adaptive hypothermic response compatible organ function.

To explore whether HGP controls insulinemia in response to infection, we used *G6Pc1^Alb11/11^* mice lacking hepatic glucose-6-phophatase c 1(G6pc1)^62^, which do not develop infection- triggered hyperglycemia (*Extended Data Fig. 17A*)^21^. Glycemia and insulinemia were lower in infected *G6Pc1^Alb11/11^ vs. G6Pc1^fl/fl^* mice (*Extended Data Fig. 17A*). This suggests that HGP contributed critically to the control of insulinemia in response to infection.

While BAT thermogenesis was not significantly reduced, body temperature was lower in infected *G6Pc1^Alb11/11^ vs. G6Pc1^fl/fl^* mice (*Extended Data Fig. 17B*). Heat dissipation was also increased in infected *G6Pc1^Alb11/11^ vs. G6Pc1^fl/fl^* mice (*Extended Data Fig. 17B*) suggesting that the adaptive thermoregulatory response to infection is partially HGP-dependent.

Adipocyte lipolysis was enhanced in infected *G6Pc1^Alb11/11^ vs.* control *G6Pc1^fl/fl^* mice, as suggested by HSL activation, but not ATGL expression, in eWAT (*Extended Data Fig. 18A - C*). This is consistent with the lower insulinemia of infected *G6Pc1^Alb11/11^* mice having a reduced capacity to repress adipocyte lipolysis^28^.

We then asked whether the regulation of insulinemia via adipocyte lipolysis^56^ regulates insulinemia in response to infection. In support of this hypothesis, the reduction of insulinemia was more pronounced in infected *Pnpla2^AdipoQ11/11^ vs. Pnpla2^fl/fl^* mice (*Extended Data Fig. 18D*). The observation that insulin administration reduced glycemia to a similar extent in infected- *Pnpla2^AdipoQ11/11^ vs. Pnpla2^fl/fl^* mice (*Extended Data Fig. 18E*), suggests that adipocyte lipolysis supports insulinemia via insulin production rather than insulin resistance.

Finally, we asked whether the protective effect of insulin against infection is mediated via INSR signaling in adipocytes^28,56^ and tested this hypothesis in *Insr^AdipoQ11/11^* mice, carrying a deletion of the *InsR* gene specifically in adipocytes^63^. *Insr^AdipoQ11/11^* mice subjected to CLP (LD_40_) reduce body temperature more severely, *albeit* not statistically significant compared to control *Insr^fl/fl^* mice (*Fig. 4F*). This was associated with an increased in lethality compared to control *Insr^fl/fl^* mice (*Fig. 4F*) and suggests that the protective effect of insulin against sepsis acts via INSR signaling in adipocytes. Considering that insulin inhibits lipolysis^28^, we infer that INSR signalling in adipocytes “guards” the extent of WAT lipolysis in response to infection, while supporting BAT thermogenesis.

Our findings demonstrate that the induction of adipocyte lipolysis is an evolutionary conserved defense strategy against infection. It is required to rewire host energy allocation^42^ and prevent life-threatening hypothermia from compromising organ function in response to infection, therefore preventing the onset of sepsis^1^. This defense strategy does not interfere with pathogen burden (*Fig. 1G, Extended Data 8B, 15E*), establishing disease tolerance to infection.

Adipocyte lipolysis does not appear to exert a major effect on the systemic inflammatory response to infection (*Extended Data Fig. 12*), therefore presumably acting downstream from the inflammatory cytokines driving the pathogenesis of sepsis^16,64–66^. This is consistent with the unfettered inflammatory response to infection dysregulating the adaptive metabolic response to infection^67^.

Our findings suggest that adaptive thermoregulation is also an evolutionary conserved adaptive response to infection^68^. While humans typically develop a hyperthermic response to infection, a fraction of sepsis patients develops hypothermia^69^ and become at the highest risk of mortality^70,71^. Moreover, therapeutic hyperthermia improves sepsis survival in afebrile sepsis patient^72^, suggesting that life-threatening hypothermia represents an evolutionary trade-off^73^ from an adaptive hypothermic response to infection^67,74^.

The strict reliance on adipocyte lipolysis to sustain the adaptive hypothermic response to infection (*Fig. 3A,B*) is reminiscent of the reliance of torpor on adipocyte lipolysis^75^. This is consistent with adipocyte lipolysis supplying FA to fuel mitochondrial fatty acid-oxidation^42^ and “guard” a lower threshold of body temperature compatible with organ function in response to infection^7,9,76^. Whether sepsis patients that develop hypothermia fail to mobilize FA via adipocyte lipolysis remains however to be established.

The sustained hypometabolic response of infected mice is preceded by a transient hypermetabolic response supported by the induction of insulin production (*Fig. 4A*) in response to HGP (*Extended Data Fig. 17A*) and adipocyte lipolysis (*Extended Data Fig. 18D*). Pharmacologic ablation of pancreatic β-cells to suppress insulin production causes a profound dysregulation of the hypothermic response to infection (*Fig. 4C,D*), compromising survival in sepsis (*Fig. 4D*). This suggests that insulin is essential to “guard” the lower threshold of the adaptive hypothermic response to infection (*Fig. 4D*). The mechanism underlying this thermoregulatory effect of insulin operates most likely via INSR signaling in different cellular compartments partaking in the adaptive hypometabolic response to infection, including in adipocytes (*Fig. 4F*). This is consistent with INSR signaling promoting cellular FA uptake to support non-shivering BAT thermogenesis^56^. Of note insulin represses adipocyte lipolysis while promoting *de novo* lipogenesis^28^, which is likely important to prevent sepsis-associated cachexia, a life-threatening condition characterized by the loss of fat and lean mass^77,78^. To what extent insulin exerts a similar salutary effect that justifies its widespread use in sepsis patients is unclear^32^.

The widespread notion that INSR signaling^28^ represses HGP^29^ to sustain an adaptive hypoglycemic response to bacterial infection^22^ is not supported by the observation that pharmacologic ablation of pancreatic β-cells does not impair the development of hypoglycemia in infected mice (*Extended Data Fig. 16C*). This suggests that the adaptive hypoglycemic response to infection operates via a mechanism that is not mediated by insulin.

In conclusion, adipocyte lipolysis plays a central role in the regulation of the hypometabolic response that establishes disease tolerance to infection and prevents the onset of sepsis. This metabolism-based defense strategy acts downstream of the inflammatory response to infection and provides an array of novel therapeutic strategies towards the treatment of sepsis.

## Material and Methods

### Animals

Mice (male and female) were bred and maintained under specific pathogen-free (SPF) conditions at the Gulbenkian Institute for Molecular Medicine, housed at standard *vivarium* temperature (21-23°C) or when indicated at 26-27°C, in a12-hour light/dark cycle with free access to water and standard chow pellets. Experimental protocols were approved by the Gulbenkian Institute for Molecular Medicine Ethics Committee/ORBEA and the Portuguese National Entity (Direção-Geral de Alimentação e Veterinária). Experimental procedures were performed according to the Portuguese legislation on protection of animals and European (Directive 2010/63/EU) legislations. C57BL/6J *AdipoQ^Cre/Wt^Pnpla2^fl/fl^* (*i.e., Pnpla2^AdipoQ^*^11/11^) mice and littermate control *Pnpla2^fl/fl^* mice were previously described^44^. C57BL/6J *Sa^CreERT2/Wt^G6pc1^fl/fl^* (*i.e., G6pc1^Alb^*^11/11^) mice and littermate control *G6pc1^fl/fl^* mice were previously described^62^. Conditional deletion of the *G6pc1^fl/fl^* allele in hepatocytes was induced by Tamoxifen (TAM; Sigma) administration starting at 8-week after birth (i.p., 10 mg/mL, 100µL, daily for 5 days), as described^62^. *G6pc1^Alb^*^11/11^ mice were used within 2 weeks after the last TAM administration, as described^21^. C57BL/6J *Insr^AdipoQ11/11^* mice carrying a deletion of the INSR specifically in adipocytes and littermate control *Insr^fl/fl^* mice were previously described^63^. Streptozocin (STZ; Abcam Cat# ab142155) was administered (i.p., 100 µg/g diluted in PBS, in 2 consecutive days) to ablate pancreatic β-cells, as described^79^. Mice were used 3 weeks after the last STZ injection. Control mice received vehicle (PBS; i.p., 10 µL/g, i.p.)

### Human adipose tissue samples

Human visceral WAT samples were obtained from patients undergoing therapeutic laparotomy as part of the INSIGHT study^49^. Peritoneal sepsis was defined according to the German sepsis guidelines^80^. Control subjects were defined as those without sepsis and without the diagnosis of type 2 diabetes mellitus or metabolic syndrome, according to the American Diabetes Association (ADA) 2010 criteria for the diagnosis of type 2 Diabetes (T2D) or the NCEP-ATPIII criteria, respectively (NCEP Expert Panel 2002; ADA 2010). Visceral adipose tissue samples were taken from the *greater omentum* immediately after midline incision and adhesiolysis and fixed in 4.5% formaldehyde (Roti Histofix 4.5%, Carl Roth, Germany) at 4°C for 24 h. Tissues were washed in double distilled water (1 hour) before dehydration in an ascending ethanol and xylene series (Carl Roth, Germany) overnight, embedded in paraffin and sectioned at 3 µm using a rotary microtome (Leica, Germany). The INSIGHT study is registered in the German Clinical Trials Register (DRKS00005450) and was approved by the faculty’s ethics review board of Jena University Hospital (3247-09/11). All subjects or their legal representatives gave written informed consent.

### Experimental polymicrobial infection

Mice were subjected to cecal ligation and puncture (CLP), essentially as described^21^. Surgeries were performed between 7:30-9:30 AM and CLP was aimed at a mortality rate of ∼20-50% in control animals (LD_20-50_). The cecum was ligated, double-punctured (23 Gauge needle), cecum content was extruded, and the cecum was placed back into the abdominal cavity. Mice received 0.9% saline (40 mL/kg, i.p.) immediately after CLP. For the peritoneal contamination and infection (PCI) model gut content was collected from Rag2^-/-^ mice and essentially prepared and conserved as described^81^. PCI suspension was injected *i.p.* (4.7-5 µL/g mouse). Mice were treated with Imipenem/Cilastatin (25 mg/kg, i.p. in the morning, 25mg/kg or 50mg/kg, i.p. in the evening) 2h after CLP/PCI and every 12h for 3 days. Mice were individually caged after CLP/PCI, except on Fig. 1G and 3C. Survival and disease severity score (DSS, including activity, level of consciousness and response to stimulus) were assessed twice daily for the first 3 days and once daily for 7-14 days. Temperature (Rodent Thermometer BIO-TK8851, Bioseb, France) was monitored twice daily for the first 3 days and once daily for 7-14 days. Weight was monitored daily. Blood glucose levels (ACCU-CHECK Aviva, Roche, Amadora, Portugal) from tail vein was monitored twice daily for the first 3 days and once daily for 7-14 days.

### Food intake and pair feeding

Steady state daily food intake was quantified for 1 day before CLP, and subsequently throughout the course of infection, essentially as described^43^. Briefly, C57BL/6J mice were housed in individual cages with free access to water and standard chow pellets. Food pellets were weighed (Ohaus^®^ CS200 scaler, Sigma Aldrich) daily at 9:00-10 AM. Daily food intake was calculated as follows: Pellet Mass_24hrs before_ - Pellet Mass _24hrs after_. Mice subjected to CLP were fed *ad lib* and food intake was determined. Pair-feed control mice underwent sham surgery. The average food intake of CLP mice was determined, and weight matched diet was given to the pair-fed control mice^82^.

### Adipose tissue mass & morphometric analyzes

Mice were euthanized, perfused *in toto* with ice-cold PBS and eWAT, scWAT and BAT were dissected and weighed (Entris^®^ II, Satorius). Tissues were fixed (10% formalin), embedded in paraffin, sectioned (3μm) and stained with Hematoxylin & Eosin (H&E). Whole sections were analyzed and images acquired with a Leica DMLB2 microscope (Leica) and NanoZoomer-SQ Digital slide scanner (Hamamatsu). For WAT adipocytes area and BAT lipid droplets measurements tissues were processed to paraffin-embedded sections (3 μm sections), stained with H&E and scanned into digital images (NanoZoomer-SQ Digital slide scanner-Hamamatsu). The average WAT adipocyte size in adipose tissue sections (expressed as the average cross-sectional area per cell [μm^2^]) was determined using Fiji software, as described^83^. Briefly, a slide scanned picture was captured at 2.5x. An average of 1500 adipocytes were measured per sample. The following macro was applied: run (“Set Scale…”, “distance = 560 known = 250 pixel = 1 unit = um global”); run (“Duplicate…”, “ ”); run (“Subtract Background…”, “rolling = 50 light separate sliding”); run (“Despeckle”); run (“8-bit”); setAutoThreshold(“Mean dark”);//run (“Threshold…”);//setThreshold(250, 255); setOption(“BlackBackground”, false); run (“Convert to Mask”); run (“Make Binary”); run (“Dilate”); run (“Close-”); run (“Invert”); run (“Analyze Particles…”, “size = 330–15000 circularity = 0.50–1.00 display exclude clear summarize add”). The average BAT lipid droplet size [expressed as the average cross-sectional area per lipid droplet (μm^2^)], and the number of lipid droplets were quantified in 3 non-overlapping 40x fields for each mouse. An average of 11000 lipid droplets per mouse were measured. The following macro was applied: run (“Set Scale…”, “distance = 452 known = 100 pixel = 1 unit = µm global”); run (“Duplicate…”, “ ”); run (“Subtract Background…”, “rolling = 50 light separate sliding”); run (“Despeckle”); run (“8-bit”); setAutoThreshold(“Mean dark”);//run (“Threshold…”);//setThreshold(245, 255); setOption(“BlackBackground”, false); run (“Convert to Mask”); run (“Make Binary”); run (“Dilate”); run (“Close-"); run (“Invert”); run (“Watershed”); run (“Analyze Particles…”, “size = 1–5000 circularity = 0.4–1.00 display exclude clear summarize add”).

### Untargeted Lipidomics

Blood was collected through intracardial puncture (23 G needle pre-coated with 0.5M EDTA), plasma was extracted by centrifugation (12.000xg; 10 min) and Plasma (50 µL) used for lipid extraction. Isopropanol (150 µL containing internal standards, SPLASH Lipidomix; Avanti Polar Lipids, AL, USA) was added and after thorough vortexing and incubation (-20 °C; 20 min) samples were centrifuged (10 min; 15,000*g*, 4 °C). Supernatants were transferred to analytical glass vials for LC-MS/MS analysis, performed on a Vanquish UHPLC system coupled to an Orbitrap Exploris 240 high-resolution mass spectrometer (Thermo Scientific, MA, USA) in negative and positive ESI (electrospray ionization) mode. Chromatographic separation was carried out on an ACQUITY Premier CSH C18 column (Waters; 2.1 mm x 100 mm, 1.7 µm) at a flow rate of 0.3 mL/min The mobile phase consisted of water:ACN (40:60, v/v; mobile phase A) and IPA:ACN (9:1, v/v; mobile phase B), which were modified with a total buffer concentration of 10 mM ammonium acetate + 0.1 % acetic acid (negative mode) and 10 mM ammonium formate + 0.1% formic acid (positive mode), respectively. The following gradient (23 min total run time including re-equilibration) was applied (min/%B): 0/15, 2.5/30, 3.2/48, 15/82, 17.5/99, 19.5/99, 20/15, 23/15. Column temperature was maintained at 65°C, the autosampler was set to 4°C and sample injection volume was set to 3 µL (positive mode) and 5 µL (negative mode). Analytes were recorded via a full scan with a mass resolving power of 120,000 over a mass range from 200 – 1700 *m/z* (scan time: 100 ms, RF lens: 70%). To obtain MS/MS fragment spectra, data-dependent acquisition was carried out (resolving power: 15,000; scan time: 54 ms; stepped collision energies [%]: 25/35/50; cycle time: 600 ms). Ion source parameters were set to the following values: spray voltage: 3250 V / 3000 V, sheath gas: 45 psi, auxiliary gas: 15 psi, sweep gas: 0 psi, ion transfer tube temperature: 300°C, vaporizer temperature: 275°C. Level 1 feature identification^44^ was based on the MS-DIAL LipidBlast V68 library.

### Confocal microscopy

Mice were sacrificed and perfused (20mL, ice-cold PBS, 0.2 mM EDTA). The adipose tissues were harvested, fixed (4% paraformaldehyde, PBS, overnight), washed (PBS, 3x, 30 min each) and permeabilized (B1N buffer: 0.1% Triton X-100, 0.3 M Glycine, 0.01% Sodium Azide in distilled water, pH 7) overnight. Samples were blocked (PtxwH buffer; 1x PBS, 0.1% Triton X-100, 0.05% Tween-20, 2mg/ml Heparin, 0.01% Sodium Azide in distilled water, 3% donkey serum) overnight at room temperature (RT). Samples were incubated with anti-ATGL (PNPLA2/ATGL Antibody, Novus biologicals, Cat#: AF5365, 1:800) and anti-pHSL (Phospho-HSL Ser660, Cat#16745804S, Cell Signaling Technology, 1:600) primary antibodies, in blocking buffer (3 days, RT; shaking) followed by washing (PtxwH buffer, 5x 1 hour each). Samples were incubated with secondary antibody, Donkey anti-Sheep IgG H&L (Alexa Fluor® 647, 1:400; Abcam, Cat#ab150179) and Donkey Anti-Rabbit IgG H&L (Alexa Fluor® 568, 1:400; Abcam, Cat#ab175470) in blocking buffer (overnight, RT, shaking) followed by washing (PtxwH buffer 5x, 30 min. each). Samples were incubated (2hrs., 37°C) with BODIPY 493/503 (Cayman) and DAPI (in PBS) and washed (PBS, 3x, 30 min). Samples were imaged using a laser scanning confocal microscope (LSM980, Zeiss) and fluorescence intensity was measured using ImageJ 1.53k software (Rasband, W.S., ImageJ, U.S. NIH, Bethesda, Maryland, USA, https://imagej.nih.gov/ij/,1997-2014).

### Human visceral adipose tissue paraffin sections

Sections were deparaffinized and incubated (90 min) in citrate buffer (10mM Sodium Citrate, 0.05% Tween 20, pH 6.0) at 99°C to retrieve antigens. After permeabilization, samples were blocked (3% donkey serum in PBS- Tween20 0.05%). Samples were then incubated with primary antibodies, ATGL (PNPLA2/ATGL Antibody, Novus biologicals, Cat#AF5365, 1:50), and pHSL (Phospho-HSL (Ser660) Antibody, Cat#16745804S, Cell Signaling Technology, 1:50) in blocking buffer (overnight). After washing (4x, PBS-Tween20 0.05%) samples were incubated with secondary antibody, Donkey Anti-Sheep IgG H&L (Alexa Fluor® 647, Abcam, CAT#ab150179, 1:100) and Donkey Anti-Rabbit IgG H&L (Alexa Fluor® 568, Abcam, Cat#ab175470, 1:100), in blocking buffer for 2 hrs. After washing (4x, PBS-Tween20 0.05%) samples were incubated (30 min) with DAPI (in PBS-Tween20 0.05%). After washing (4x, PBS-Tween20 0.05%) samples were mounted with Fluoromount-G™ Mounting Medium (Invitrogen, Cat#00-4958- 02) and imaged using a laser scanning confocal microscope (LSM980, Zeiss). Fluorescence intensity was measured using ImageJ 1.53k software (Rasband, W.S., ImageJ, U.S. NIH, Bethesda, Maryland, USA, https://imagej.nih.gov/ij/,1997-2014).

### Serology

Mice were euthanized and blood collected through intracardial puncture (23 G needle pre-coated with 0.5M EDTA). Serological analysis was performed by DNAtech (Portugal; http://www.dnatech.pt/web/).

### Cytokine Measurements

Mouse cytokines were measured using the LegendPlex^TM^ Mouse Inflammation Panel (13-plex, BioLegend, Cat#740446).

### Insulin Measurements

Mouse insulin was quantified in plasma by ELISA (Crystal Chem; Cat#90080).

### Western Blot

Mice were euthanized, perfused (20 mL, ice-cold PBS + 0.2 mM EDTA), BAT was collected, snap frozen in liquid nitrogen and homogenized in NP40 extraction buffer (100mL of extraction buffer contain 3mL of 5M NaCl, 10mL of 10% NP-40, 5mL of 1M Tris, 82mL of H2O, protease inhibitor: complete Mini, EDTA-free, Sigma Cat#11836170001) with a tissue lyser (Qiagen) with tungsten carbide beads (Qiagen). Homogenate was transferred and equal amount of NP40 extraction buffer was added. Samples were incubated (1hrs on ice) and centrifuged (4°C, 10min, 20.000xg). Supernatant was transferred and protein was quantified by Bradford assay (BioRad, Cat#5000006). Protein (30 µg) was resolved on a 10% SDS-PAGE and transferred to Polyvinylidene fluoride (PVDF) membranes. The membranes were blocked (5% bovine serum albumin: BSA in 1XT-TBS) and incubated with primary antibodies overnight (4°C): Anti-mouse phospho-AKT (Ser473) (Cat#4060, Rabbit, 1:1000), Anti-mouse AKT (Cat#9272, Rabbit, 1:1000) and anti-β-Actin (Cat#4967, Rabbit, 1:5000) all from Cell Signaling Technology. Membranes were washed (3x, 1XT-TBS) and incubated (2hrs, RT) with corresponding peroxidase-conjugated secondary antibodies (Goat anti-Rabbit IgG (H+L) HRP #31460. Peroxidase activity was detected using SuperSignal West Pico PLUS Chemiluminescent Substrate (ThermoFisher Scientific). Blots were developed using AmershamImager 680 (GE Healthcare), equipped with a Peltier cooled Fujifilm Super CCD. Densitometry analysis was performed with ImageJ (Rasband, W.S., ImageJ, U.S. NIH, Bethesda, Maryland, USA, https://imagej.nih.gov/ij/, 1997-2014), using only images without saturated pixels.

### Histology (Heart, Liver, Kidney)

Mice were sacrificed, perfused *in toto* (20 mL ice-cold PBS + 2mM EDTA), organs were harvested, fixed in formalin (10%), embedded in paraffin, sectioned (3µm) and stained with Hematoxylin & Eosin (H&E). Whole-slide images were scanned using the NanoZoomer-SQ Digital Slide Scanner (Hamamatsu Photonics) and analyzed and captured with the NDP.view2 software (Hamamatsu Photonics).

### Pathogen Load

Peritoneal fluid was obtained by peritoneal lavage (7 mL sterile PBS). Mice were perfused (20 mL ice-cold PBS), whole organs were harvested and homogenized under sterile conditions in 1 mL sterile ice-cold PBS using a dounce tissue grinder (Sigma Cat#D8939-1SET). Serial dilutions were plated onto TrypticaseSoy Agar II with 5% Sheep Blood plates (Becton Dickinson Cat#254053) and incubated (24hrs, 37°C) in air 5% CO_2_ (aerobes) or in an airtight container equipped with the GasPak anaerobe container system (Becton Dickinson Ref 260678). Anaerobic conditions were confirmed in all experiments using BBL Dry Anaerobic Indicator Strips (Becton Dickinson Cat#271051).

### Infrared Temperature Measurements

At least 24h prior the measurements, mice were anesthetized (1-2% Isoflurane) and the interscapular area was shaved. BAT and tail temperatures were measured in mice that were allowed to move freely in a cage, using an infrared camera (FLIR E96: Compact-Infrared-Thermal-Imaging-Camera; FLIR Systems; West Malling, Kent, UK). Images were analyzed with a software package (FLIR Tools^®^ Software, FLIR Systems; West Malling, Kent, UK), as described^83^. Individual BAT and tail temperatures were calculated from 2-4 images taken per mouse, as per manufacturer recommendations (FLIR Tools Software, FLIR Systems; West Malling, Kent, UK), as described^83^.

### Promethion behavioral and phenotyping system

Measurements of indirect calorimetry were performed using Promethion Core (Sable Systems, USA). Mice were housed on a 12/12h light/dark cycle with controlled temperature and humidity. After 1-2 days of acclimation phase, CLP was performed. The recording continued for the following 4 days. The system is equipped with a standard GM-500 cage, with a food hopper and a water bottle connected to load cells (2 mg precision) with 1 Hz rate data collection. Additionally, the cage contains a red house enrichment. Ambulatory activity was monitored at 1 Hz rate using an XY beam break array (1 cm spacing). Oxygen, carbon dioxide and water vapor were measured using a CGF unit (Sable Systems). This multiplexed system operated in pull-mode. Air flow was measured and controlled by the CGF (Sable Systems) with a set flow rate of 2 L/min Oxygen consumption and carbon dioxide production were reported in milliliters per minute (mL/min). Energy expenditure was calculated using the Weir equation^84^ and Respiratory Exchange Ratio (RER) was calculated as the ratio of VCO_2_/VO_2_. Raw data was processed using Macro Interpreter v2.41(Sable Systems).

### Untargeted Metabolomics. Sample preparation (Heart, Brain, Liver)

Organs were harvested after perfusion (20 mL ice-cold PBS), snap frozen in liquid nitrogen and stored at - 80°C until further processing. Extraction of polar metabolites was conducted via a biphasic extraction procedure. Initially, a corresponding volume of 80% MeOH (containing internal standards, ^13^C and ^15^N labelled amino acids) was added to each sample to yield a uniform concentration of 50 mg/mL. Samples were then homogenized on dry ice via a bead beater (FastPrep-24; MP Biomedicals, CA, USA) at 6.0 m/s (5 x 30 s, 5 min pause time) using 1.0 mm zirconia/glass beads (Biospec Products, OK, USA). Subsequently homogenized suspension (200 µL) was transferred to a new sample tube and MTBE (500 µL) was added. The monophasic mixture was vortexed for 60 s and incubated (-20°C, 20 min). For phase separation, water was added (125 µL), followed by another vortexing and incubation step (see previous conditions). The biphasic solvent system was then centrifuged (10 min, 15,000xg, 4 °C, 5415R microcentrifuge (Eppendorf, Hamburg, Germany)). Afterwards, the bottom aqueous phase (220 µL) was transferred, dried under a stream of nitrogen, and reconstituted (100 µL, 80% MeOH v/v). The final samples were vortexed for 10 min, centrifuged (see previous conditions) and the supernatants were transferred to analytical glass vials for LC-MS/MS analysis.

### Untargeted Metabolomics. LC-MS/MS analysis

LC-MS/MS analysis was performed on a Vanquish UHPLC system coupled to an Orbitrap Exploris 240 high-resolution mass spectrometer (Thermo Scientific, MA, USA) in negative and positive ESI (electrospray ionization) mode. Chromatographic separation was carried out on an Atlantis Premier BEH Z-HILIC column (Waters, MA, USA; 2.1 mm x 100 mm, 1.7 µm) at a flow rate of 0.25 mL/min The mobile phase consisted of water:acetonitrile (9:1, v/v; mobile phase phase A) and acetonitrile:water (9:1, v/v; mobile phase B), which were modified with a total buffer concentration of 10 mM ammonium acetate (negative mode) and 10 mM ammonium formate (positive mode), respectively. The aqueous portion of each mobile phase was pH-adjusted (negative mode: pH 9.0 via addition of ammonium hydroxide; positive mode: pH 3.0 via addition of formic acid). The following gradient (20 min total run time including re-equilibration) was applied (time [min]/%B): 0/95, 2/95, 14.5/60, 16/60, 16.5/95, 20/95. Column temperature was maintained at 40°C, the autosampler was set to 4°C and sample injection volume was set to 4 µL. To obtain MS/MS fragment spectra, data-dependent acquisition was carried out (resolving power: 15,000; scan time: 22 ms; stepped collision energies [%]: 30/50/70; cycle time: 900 ms). Ion source parameters were set to the following values: spray voltage: 4100 V (positive mode) / -3500 V (negative mode), sheath gas: 30 psi, auxiliary gas: 5 psi, sweep gas: 0 psi, ion transfer tube temperature: 350°C, vaporizer temperature: 300°C. All experimental samples were measured in a randomized manner. Pooled quality control (QC) samples were prepared by mixing equal aliquots from each processed sample. Multiple QCs were injected at the beginning of the analysis in order to equilibrate the analytical system. A QC sample was analyzed after every 5^th^ experimental sample to monitor instrument performance throughout the sequence. For determination of background signals and subsequent background subtraction, an additional processed blank sample was recorded. Data was processed using MS-DIAL4.9.221218^85^ and raw peak intensity data was normalized via total ion count of all detected analytes^86^. Level 1 feature identification^87^ was based on an in-house library for metabolomics (EMBL-MCF 2.0) using accurate mass, isotope pattern, MS/MS fragmentation, and retention time information with a minimum matching score of 80%.

### Quantification and statistical analysis

Mouse lipidomic data normalization. From lipidomic cohorts 1 (CLP *vs.* Ctrl/Sham) and 2 (*Pnpla2^AdipoQ11/11^ vs. Pnpla2^fl/fl^*mice), we identified 1,871 and 1,691 compounds, respectively. Features with a coefficient of variation (CV) greater than 30% in QC samples, a missing percentage greater than 30% in all samples, and a missing percentage equal to 100% in one of the study groups were removed. In total, 1,799 compounds from cohort 1 and 1,648 compounds from cohort 2 remained after quality control. Missing values were imputed using the random forest method from the R package missForest^88^. Data were log_2_ transformed for downstream analyses. Lipid classes are identified by mapping compound’s SMILES with ClassyFire^89^. Human lipidomic data has been previously published^10^ and includes patients from community acquired pneumonia (CAP) derived sepsis (N=168) and controls (N=48). Controls are subjects presenting at the outpatient clinics of the Amsterdam UMC location AMC without infection or other acute disease, but with comorbidities for which they receive outpatient treatment, as described^10^. We used a lipid annotations converter for large scale lipidomics and epilipidomics datasets (LipidLynxX) to align mouse and human lipid metabolites, converting different lipid annotations obtained in different MS machines/laboratories to a unified set of lipid identifiers^90^. This gave rise to 1022 unique lipid metabolites in mice and 1476 in humans. Using the unified identifiers to match mouse and human lipid metabolites we compared 816 lipid metabolites common to both mouse and humans and applied a comparative analysis to this dataset. Lipids were stratified into the following lipid classes defined in our lipidomics pipeline in terms of their generic chemical formula^91^. CER: ceramide, DG: diglycerides/diacylglycerols, FA: fatty acids, HCER; hexosylceramides, LPC: lysophosphatidylcholine, LPE: lysophosphatidylethanolamine, PC: phosphatidylcholine, PE: phosphatidylethanolamine, PI: phosphatidylinositol, SM: sphingomyelin/sphingolipid, ST: sterol, TG: triglycerides. Mouse metabolomic data (*Pnpla2^AdipoQ11/11^ vs. Pnpla2^fl/fl^* mice): We identified 157 metabolites in heart, 186 metabolites in liver and 161 metabolites in brain. Features with a coefficient of variation (CV) greater than 30% in QC samples. Only one of the repeated compounds detected by positive and negative ions was kept by the following order: [M+H]+, [M-H]-, followed by other adduct types. Data were log_2_ transformed for downstream analyses. Metabolites are first mapped to the Human Metabolome Database (HMDB) v5.0 based on SMILES and InChIKey, the unmapped compounds are further identified by ClassyFire based on SMILES and InChIKey, and the rest unmapped are manually mapped to Human Metabolome Database (HMDB)^92^ by name, chemical formula, and average molecular weight.

### Statistical analyses

Principal component analysis (PCA) was applied to examine the overall distribution of the sample data using the R package ’stats’. Two-sided T-tests were calculated using the R package ’rstatix’. Next, the ’cp4p’ R package^93^ was exploited to perform multiple testing correction using the ’st.boot’ method, with a P-value ≤ 0.05 and an FDR-adjusted p ≤ 0.25 considered as the significant thresholds. The over representation of the KEGG pathways for metabolites are identified by MetaboAnalyst 6.0^88^. Data visualization was performed using the R packages ’ggplot2’, ’ggvenn’, and ’ComplexHeatmap’^94^.

Comparisons of individual lipid levels between two groups (CLP vs. Control, *Pnpla2^AdipoQ11/11^ vs. Pnpla2^fl/fl^*, CAP vs. healthy Control) were quantified using log_2_ fold change, t-statistic and/or p-value from a t-test. Correction for multiple testing was performed using Benjamini-Hochberg (BH) procedure. A Venn-Euler plot was constructed using a BH-adjusted p-value threshold of 0.25 for mice and 0.05 for human. Correlation between lipid levels and SOFA in CAP patients (at ICU admission) was quantified using Pearson’s R, and a univariate linear model to determine p-value and the line of best fit (blue) with 95% confidence band (shaded grey).

Statistically significant differences between two experimental groups were assessed using, if not otherwise stated, a two-tailed t-test. Comparisons between more than two groups were carried out by one or two-way ANOVA with Tukey’s, Sidak’s or Dunn’s multiple-comparisons test. Survival was represented by Kaplan–Meier plots and difference between the groups was assessed using the log-rank test. Contingency analysis was performed using the Chi-square test. All statistical analyses were performed using GraphPad Prism 8 software. Differences were considered significant at a p value <0.05. ns: Not significant, p > 0.05; *p < 0.05, **p < 0.01; ***p < 0.001; ****p < 0.0001.

## Acknowledgements

The authors are indebted to Sofia Rebelo and all members of the Inflammation group (GIMM) for technical and intellectual contributions, to GIMM core facilities, namely histopathology, bioimaging, flow cytometry and rodent and to Ines Eichhorn for technical assistance in preparing microscopy sections of human adipose tissue.

## Author Contributions

MPS conceptualized the project with EJ and TWA.

EJ designed and performed all the experimental work, with initial contribution from TWA, and contributed critically to data analysis.

HPS, JB, TvdP curated and analyzed/visualized lipidomics data from human CAP patients. LLX, PF, WFGNT, AK, GP, BD curated and analyzed/visualized omics data from murine samples.

PH, UBM, CvL provided human visceral adipose tissue from septic patients. STR, JK, SP, VP, MM, SC, TWA performed experiments.

MPS wrote the manuscript with EJ.

MPS, EJ, MB, SW contributed to funding support.

MPS, SW, AM, GP, CvL, TvdP, GM supervision and project planning. MPS, EJ project administration

## Competing interests

PH reports funding from Novo Nordisk and MSD through the institution, lecture fees from Orpahalan and travel support from Falk and Ipsen. CvL received lecture fees and travel support from Fresenius-Kabi. Other authors declare no competing interests.

## Funding

This work was financed by Fundação para a Ciência e Tecnologia (2022.08590.PTDC_EXPL DOI 10.54499/2022.08590.PTDC to JK; 2023.09168.CEECIND/CP2854/CT0005 to EJ; FEDER/29411/2017, PTDC/MED-FSL/4681/2020 DOI 10.54499/PTDC/MED- FSL/4681/2020, 2022.02426.PTDC DOI 10.54499/2022.02426.PTDC, UI/BD/152257/2021 to MM and Congento LISBOA-01-0145-FEDER-022170 to MPS). DFG Cluster of Excellence ‘‘Balance of the Microverse’’ EXC 2051; 390713860 (EJ, GP, MPS as associated member). Gulbenkian Foundation (MPS and IBB 2021-51/BI-D/2021 to ST). la Caixa Foundation HR18- 00502 (EJ, JK, MPS, MB, SW). Human Frontier Science Program (LT0043/2022-L to JK). European Union’s Horizon 2020 research and innovation programme (Grant 955321). Oeiras- ERC Frontier Research Incentive Awards (MPS). H2020-WIDESPREAD-2020-5-952537 SymbNET Research Grants (MPS)

## Additional information

**Supplementary Information** is available for this paper. Extended Data Figures 1-18.

**Correspondence and requests for materials** should be addressed to and

---

**Extended Data Figure 1:**
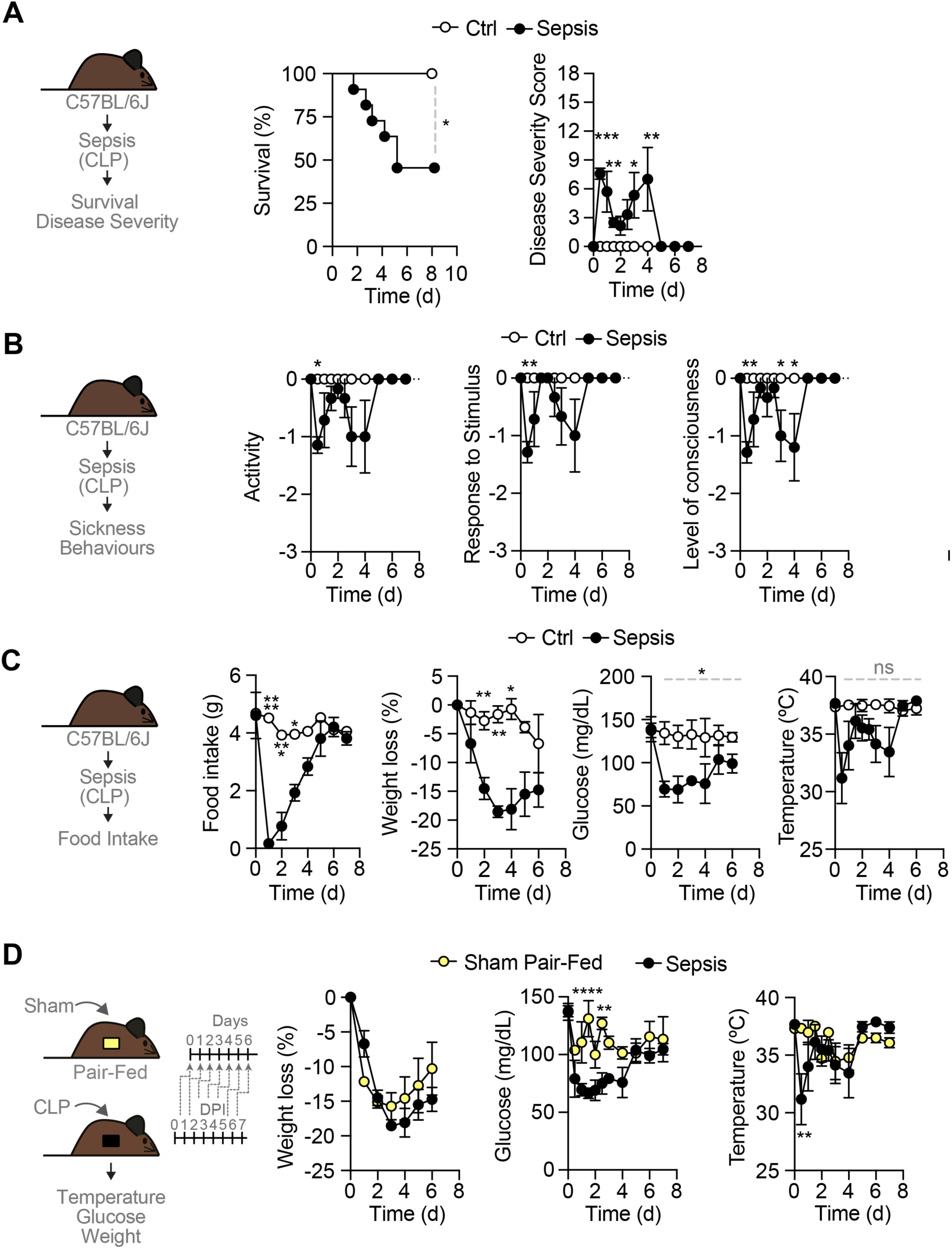
Distinct metabolic responses to infection and starvation (Related to Figure 1). **A)** Survival of mice subjected to CLP (Sepsis, N=11) *vs.* control (Ctrl, N=9) mice. Data pooled from 3 independent experiments, with similar trend. P-values calculated by Log-rank (Mantel-Cox) test, *p<0.05. Disease severity score of C57BL/6J mice subjected to CLP (Sepsis, N=7) *vs.* controls (N=5). Data represented as mean ± SD, pooled from 2 independent experiments, with similar trend. (**B**) Sickness-associated scores: activity, response to stimulus and level of consciousness of C57BL/6J mice subjected to CLP (Sepsis, N=7) *vs.* controls (N=5). Data represented as mean ± SD, pooled from 2 independent experiments, with similar trend. (**C**) Food intake, weight loss (% of initial), blood glucose, and core body temperature of C57BL/6J mice subjected to CLP (Sepsis, N=7) *vs.* control (N=5). Data represented as mean ± SD, pooled from 3 independent experiments, with similar trend. (**D**) Weight loss (% of initial), blood glucose, and core body temperature in CLP (Sepsis, N=11) *vs.* sham-operated, pair-fed C57BL/6J mice (N=12). Data represented as mean ± SEM, pooled from 3 independent experiments, with similar trend. P-values calculated by Two-Way ANOVA with Šídák’s multiple comparisons test. *p≤0.05; **p≤0.01; ***p≤0.001; ****p≤0.0001.

**Extended Data Figure 2:**
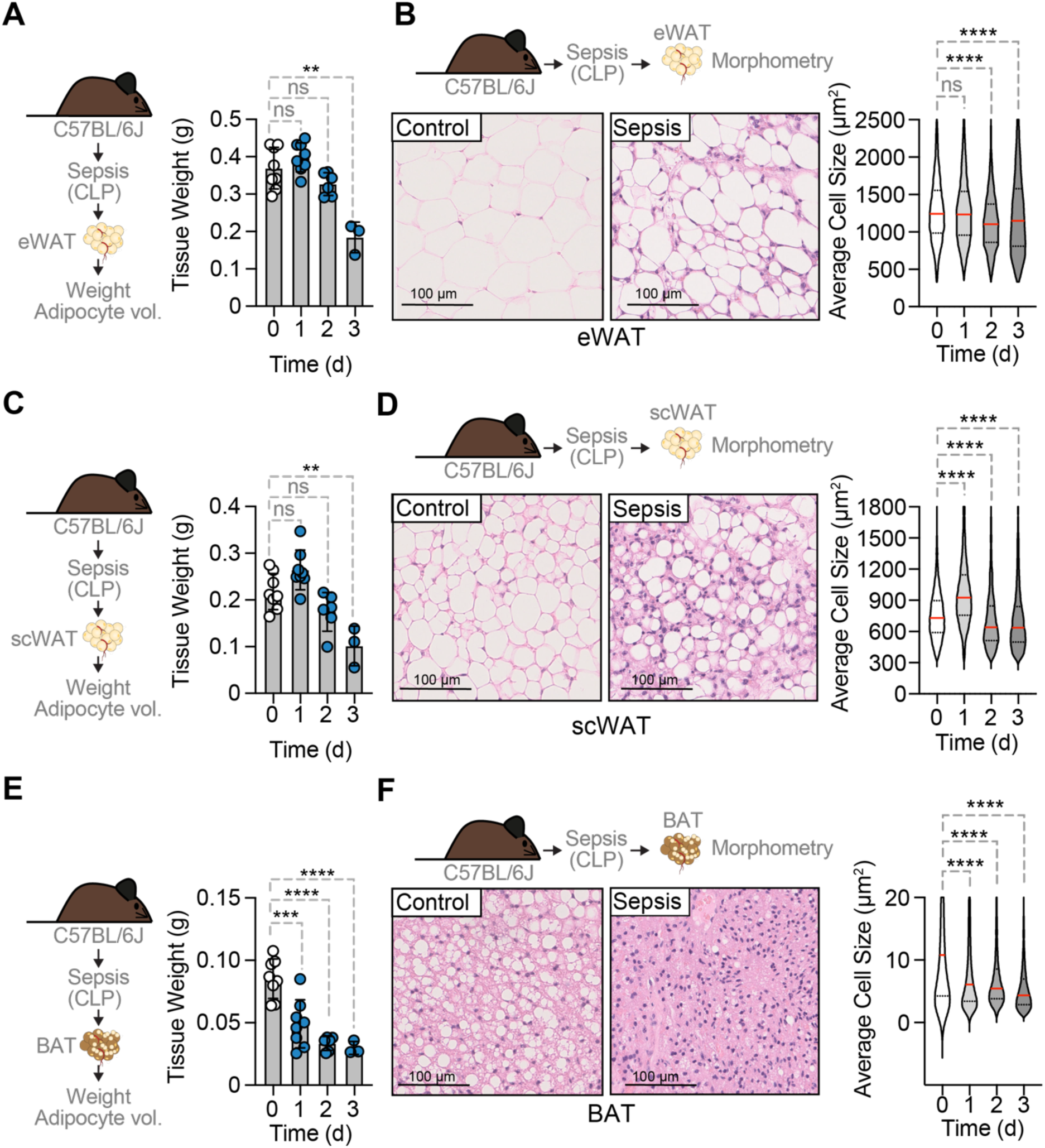
Sepsis is associated with adipose tissue wasting in mice (Related to Figure 1). **A)** Tissue weight of epididymal white adipose tissue (eWAT) in C57BL/6J mice subjected to CLP (Sepsis) *vs.* control (0d). **B)** Representative H&E staining of eWAT from CLP (Sepsis, 3d) or control (0d) C57BL/6J mice (N=3-8) and quantification of lipid vacuole size, lipid vacuole size depicted until 2500 µm^2^. **C)** Tissue weight of sub-cutaneous WAT (scWAT) in C57BL/6J mice subjected to CLP (Sepsis) *vs.* control (0d). **D)** Representative H&E staining of scWAT from CLP (Sepsis, 3d) or control (0d) C57BL/6J mice (N=3-8) and quantification of lipid vacuole size, lipid vacuole size depicted until 1800 µm^2^. **E)** Tissue weight of interscapular brown adipose tissue (BAT) weight in C57BL/6J mice subjected to CLP (Sepsis) *vs.* control (0d). **F)** Representative H&E staining of BAT from CLP (Sepsis, 3d) or control (0d) C57BL/6J mice (N=3-8) and quantification of lipid vacuole size, lipid vacuole size depicted until 20 µm^2^. A)-F) Data represented as mean ± SD, pooled from 3 independent experiments, with similar trend. Circles represent individual mice. Images are representative of 3 independent experiments. Right panel: quantification of lipid vacuole size. P-values calculated with One-Way ANOVA with Dunnett multiple comparison test, **p≤0.01; ***p≤0.001; ****p≤0.0001.

**Extended Data Figure 3:**
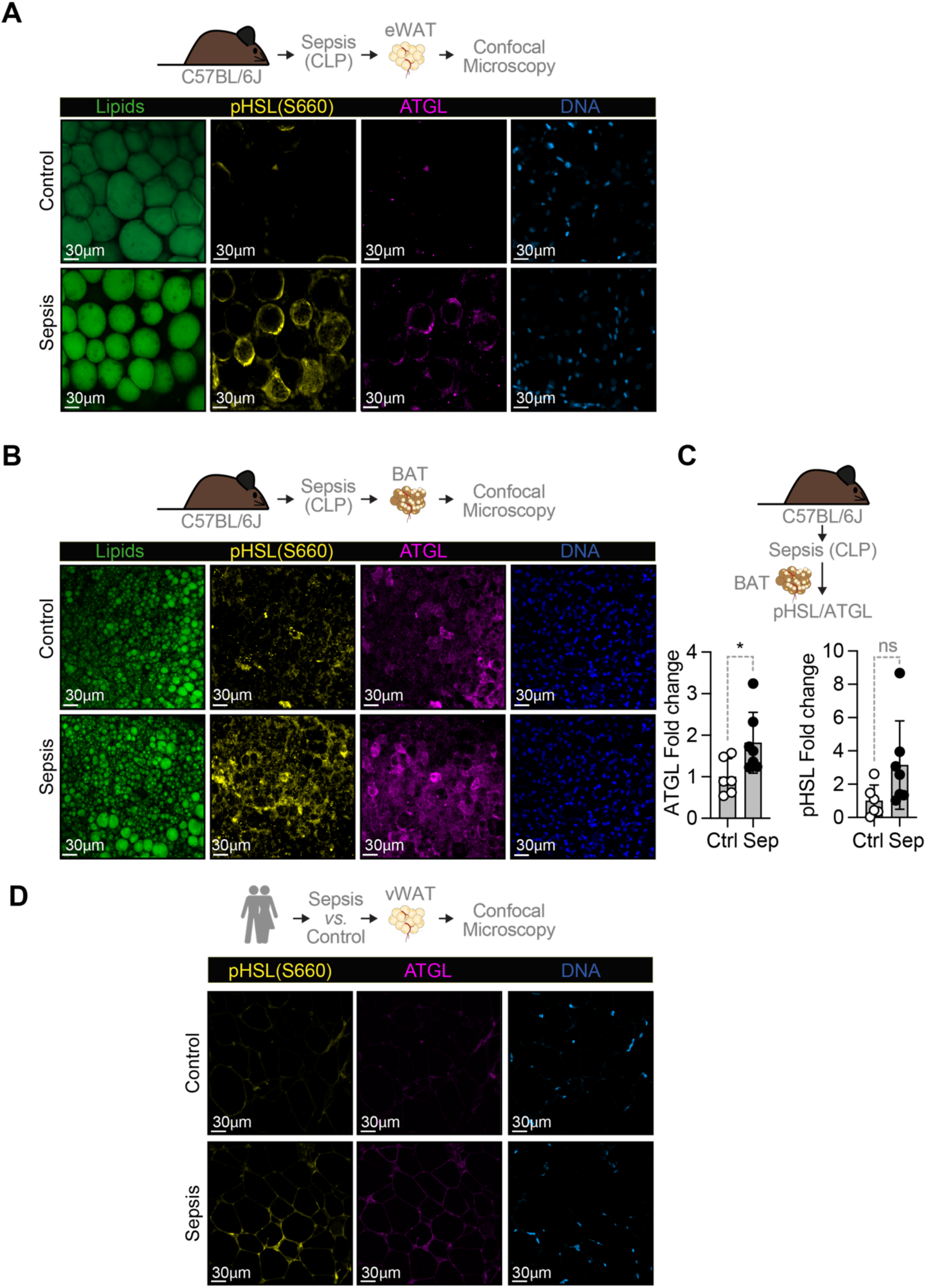
Sepsis is associated with adipocyte lipolysis in mice and humans (Related to Figure 1). **A)** Immunofluorescence imaging of neutral lipids (BodiPy; green), DNA (DAPI; blue), ATGL (anti-ATGL antibody with Alexa Fluor^®^ 647; magenta) and phospho-HSL (anti-pHSL antibody with Alexa Fluor^®^ 568; yellow) from the eWAT of C57BL/6J mice, 24hrs after CLP (Sepsis) or sham operation (Control). **B)** Immunofluorescence imaging as in A) from the BAT of C57BL/6J mice, 24hrs after CLP (Sepsis) or sham operation (Control, Ctrl). **C)** Quantification of ATGL and pHSL (N=5-7 *per* group). Data represented as mean ± SEM, from 23 different areas in the same images, pooled from 23 images in independent experiments with similar trend. Circles represent individual mice. P-values calculated by Student’s t-test, ns- not significant, *p≤0.05. **D)** Immunofluorescence imaging as in A) without neutral lipids from visceral WAT (vWAT) from septic *vs.* control individuals (N=6-7).

**Extended Data Figure 4:**
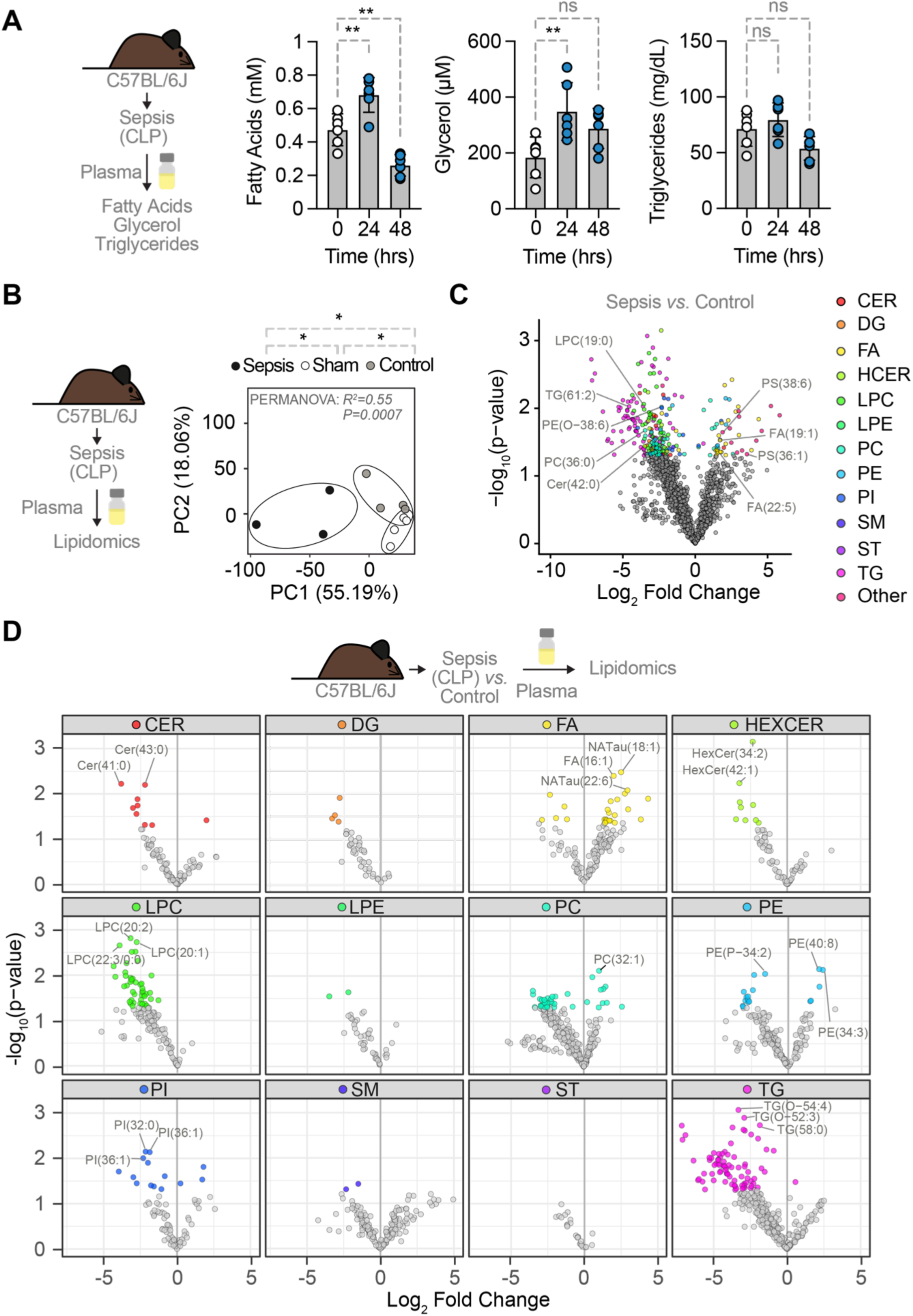
Sepsis is associated with specific plasma lipidomic profile in mice (Related to Figure 1). **A)** Plasma fatty acids, glycerol and triglycerides at different time points after CLP (Sepsis, 24hrs and 48hrs) vs. control (0hrs) in C57BL/6J mice (N=6 *per* time point). Data represented as mean ± SD. P-values calculated by One-Way ANOVA with Dunnett multiple comparison test, ns- not significant, **p≤0.01. **B)** PCA of untargeted plasma lipidomics of CLP (Sepsis, N=3, 24hrs), sham operated (N=4,24hrs) or control (N=4) C57BL/6J mice. P-values were calculated by Permanova. **C)** Differentially regulated lipid metabolites comparing sepsis (CLP) vs. control. From same mice as B). Indicated lipid metabolites are similarly regulated in human cohort (Fig. 1E,F and Fig. S5). **D)** Lipidomic profile as in C) separated by lipid class. Three most altered lipid metabolites are labeled. C)-D) Significant lipid metabolites P-value ≤ 0.05. Grey dots indicate non-significantly changed lipid metabolites.

**Extended Data Figure 5:**
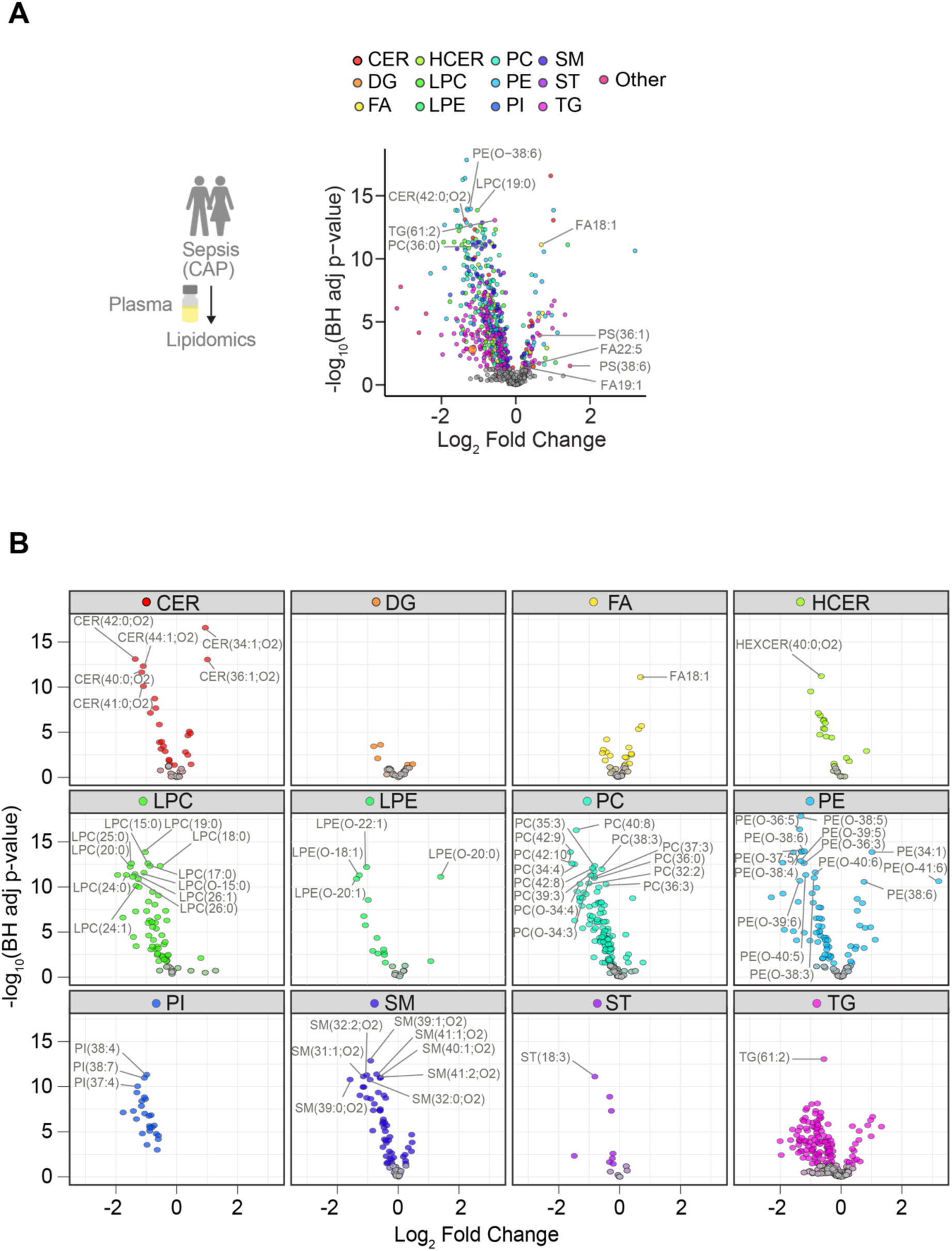
Sepsis is associated with specific plasma lipidomic profile in humans (Related to Figure 1). **A)** Differentially regulated lipid metabolites comparing septic (community acquired pneumonia (CAP) induced sepsis, N=169) *vs.* control (outpatient controls, N=48) at admission at the ICU. **B)** Lipidomic profile as in A) separated by lipid class. A)-B) Significant lipid metabolites were defined with a BH-adjusted p ≤ 0.05. Grey dots indicate non-significantly changed lipid metabolites.

**Extended Data Figure 6:**
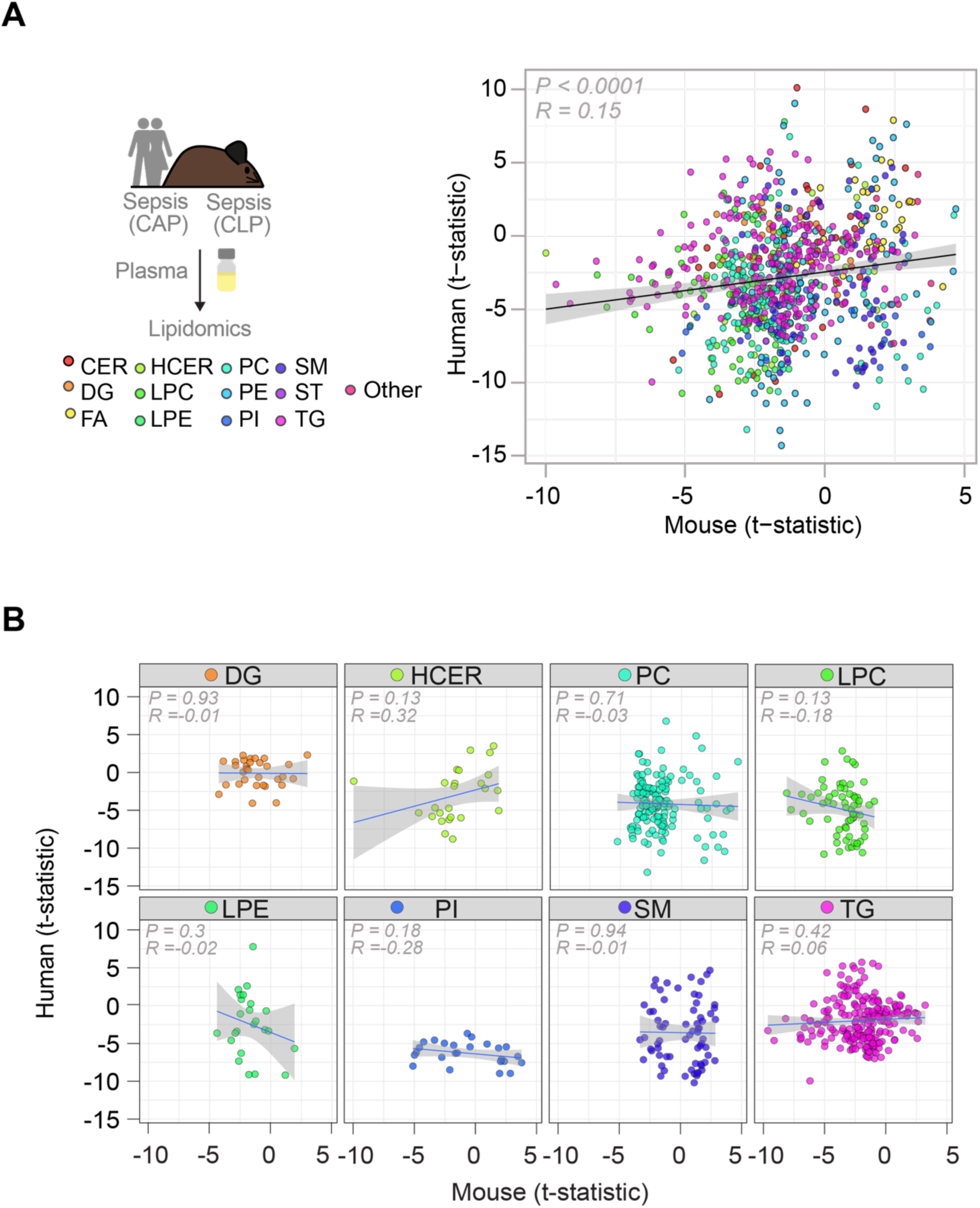
Sepsis is associated with evolutionary conserved plasma lipidomic profile in mice and humans (Related to Figure 1). **A)** Correlation between the human and mouse lipidomic response to sepsis, using the t-statistics of the plasma lipid metabolites calculated from human control (outpatient controls, N=48) *vs.* septic (community acquired pneumonia (CAP) induced sepsis, N=169) and the t-statistics calculated from control (N=4) vs. septic (CLP, N=3, 24 h) C57BL/6J mice at 24h of sepsis**. B)** As in (A) with facetted correlations of t-statistics for diacylglycerols (DG), Hexoceramides (HCER), Phosphatidylcholine (PC), Lysophosphatidylcholine (LPC), Lysophosphatidylethanolamine (LPE), Phosphatidylinositol (PI), Sphingomyelin (SM), and Triglycerides (TG).

**Extended Data Figure 7:**
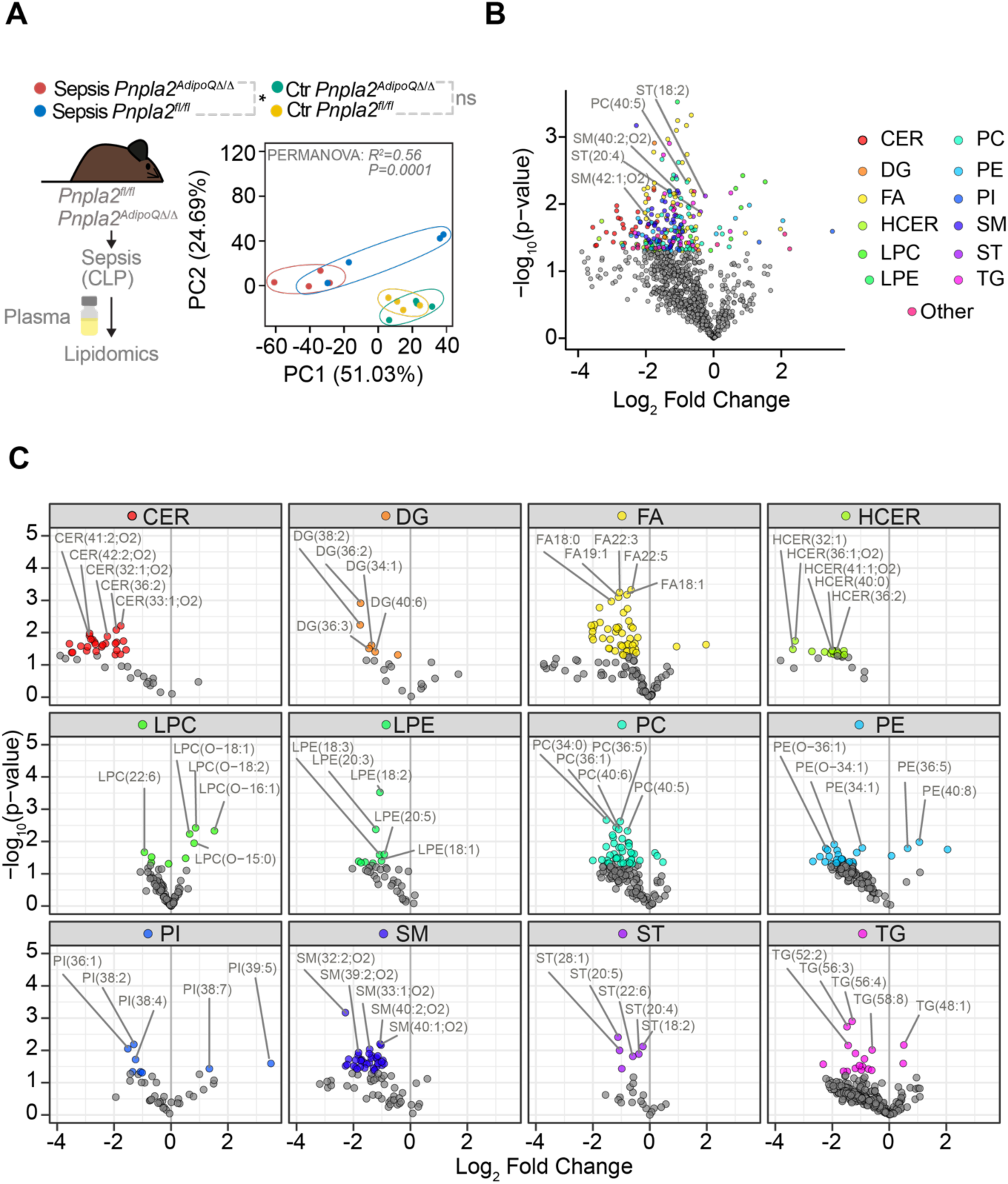
Adipocyte lipolysis determines the plasma lipidomic profile of sepsis in mice (Related to Figure 1). **A)** PCA of untargeted plasma lipidomics in *Pnpla2^fl/fl^* and *Pnpla2^AdipoQ11/11^* mice subjected to CLP (Sepsis, 24 hrs) or control. N=4 animals *per* group. **B)** Differentially regulated lipid metabolites from same mice as A) septic *Pnpla2^AdipoQ11/11^ vs*. septic *Pnpla2^fl/fl^*. **C)** Differentially regulated lipid metabolites as in B) separated by lipid class. B)-C) Significant lipid metabolites were defined with an FDR-adjusted p ≤ 0.25. Grey dots indicate non-significantly changed lipid metabolites.

**Extended Data Figure 8:**
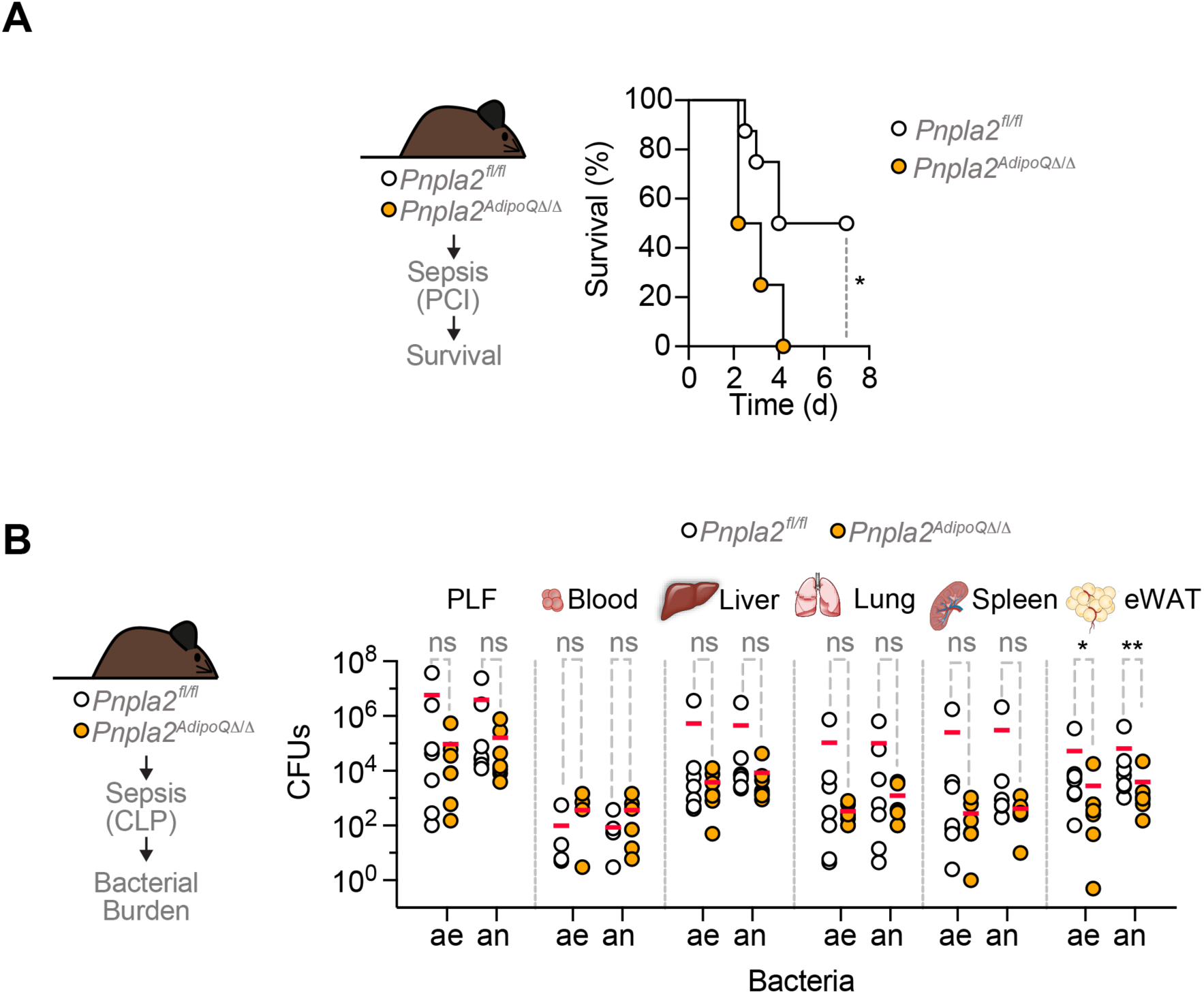
Adipocyte lipolysis establishes disease tolerance to sepsis in mice (Related to Figure 1). **A)** Survival of *Pnpla2^fl/fl^* (N=8) and *Pnpla2^AdipoQ11/11^* (N=4) mice subjected to peritoneal contamination and infection (PCI, Sepsis). Data pooled from 2 independent experiments. P-values calculated by Log-rank (Mantel-Cox) test, *p<0.05. **B)** Bacterial colony forming units (CFU; ae: aerobe, an: anaerobe) from peritoneal lavage fluid (PLF), blood, liver, lung, spleen and epidydimal white adipose tissue (eWAT) in *Pnpla2^fl/fl^* or *Pnpla2^AdipoQ11/11^* mice subjected to CLP (Sepsis, 24hrs, N=4-6). Data represented as mean (read bar), pooled from 3 independent experiments, with similar trend. Circles represent individual mice. P values calculated by Students t-test on log-transformed data, ns – not significant, *p<0.05; **p<0.01.

**Extended Data Figure 9:**
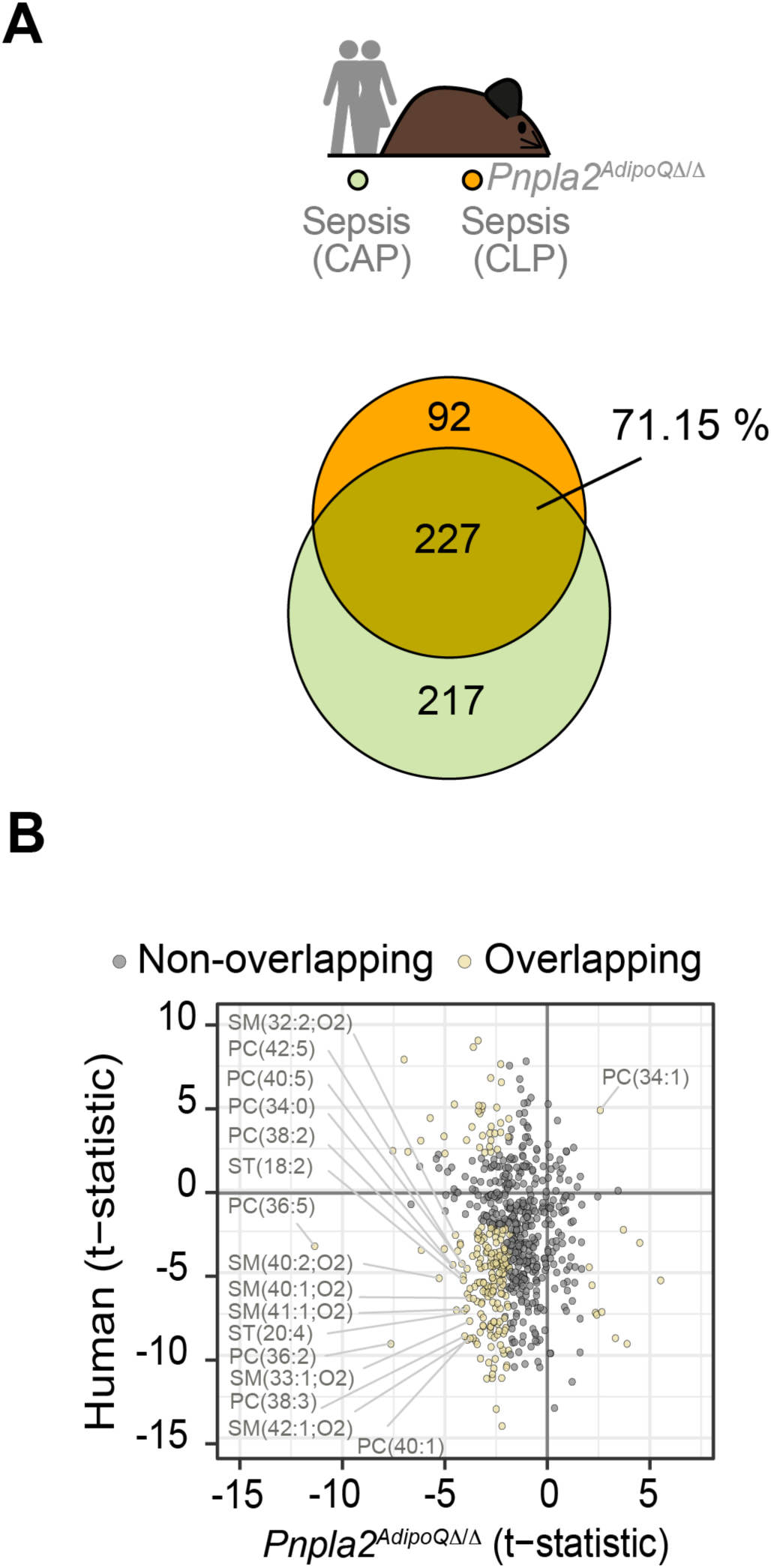
Adipocyte lipolysis determines the evolutionary conserved plasma lipidomic profile of sepsis in mice and humans (Related to Figure 1). **A)** Venn- diagram of overlapping lipid metabolites significantly differentially regulated in community acquired pneumonia (CAP) septic patients (green, N=169) and septic *Pnpla2^AdipoQ11/11^* mice (CLP, orange, N=4) *vs.* human (N=48) controls and mouse (N=4), respectively. **B)** Correlation between *Pnpla2^AdipoQ11/11^* and human lipidomic response to sepsis; using t-statistics of the plasma lipid metabolites calculated from human septic (CAP sepsis, N=169) *vs.* control (N=48) and the t-statistics calculated from septic (CLP, 24hrs) *Pnpla2^AdipoQ11/11^* (N=4) *vs. Pnpla2^fl/fl^* (N=4) mice. Yellow dots indicate the overlapping lipid metabolites between human and mice. Grey dots represent non-overlapping lipid metabolites.

**Extended Data Figure 10:**
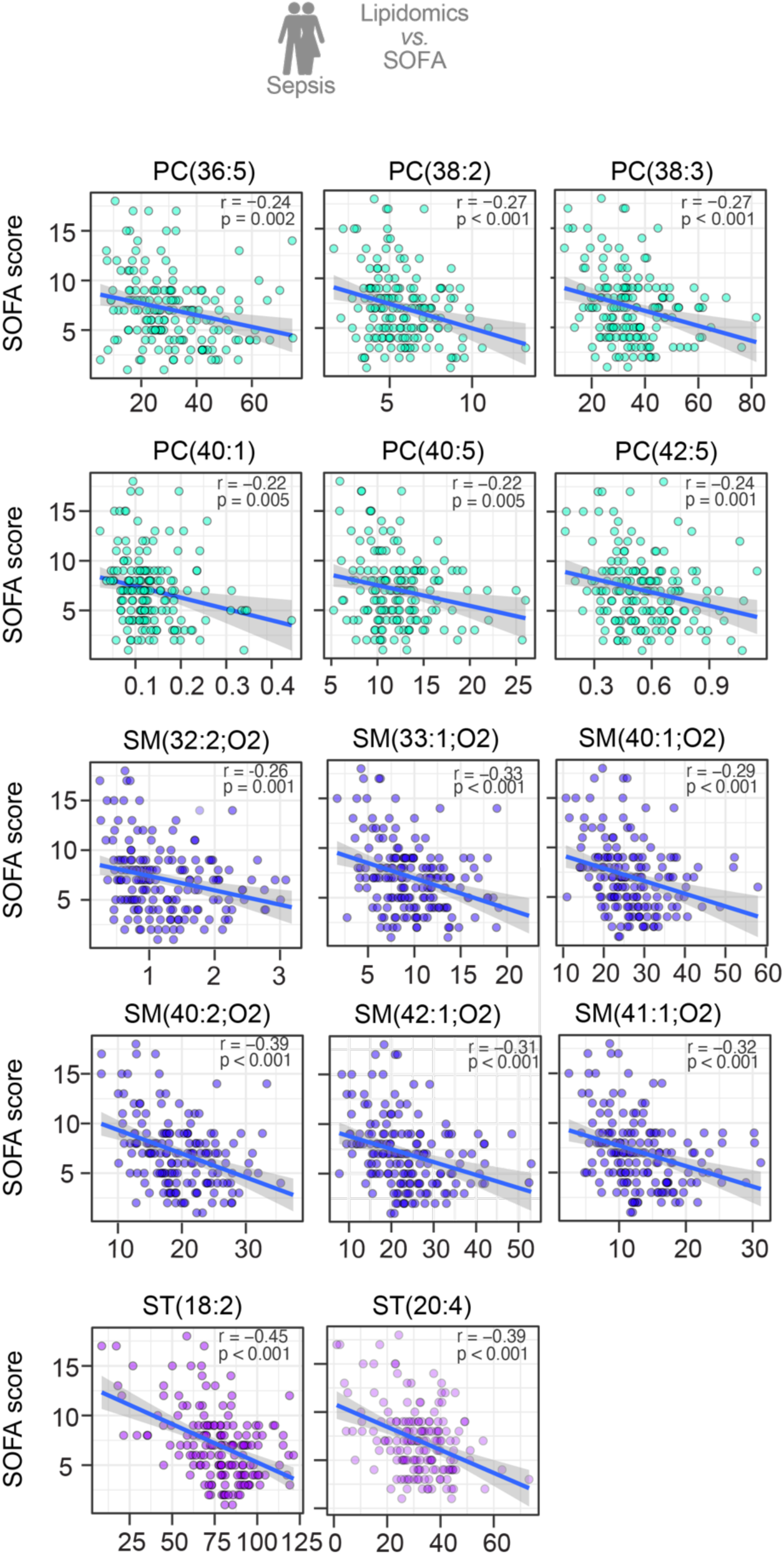
Lipid metabolites regulated via adipocyte lipolysis in infected mice, predict sepsis severity in humans (Related to Figure 1). Correlation between the sequential organ failure assessment (SOFA) score in septic (CAP) patients (N=169) and lipid levels in those patients; showing overlapping lipids that were significantly less abundant in both septic (CLP, 24hrs) *Pnpla2^AdipoQ11/11^* mice and septic (CAP) patients compared to respective controls, identified in Figure S9B (lower, left quadrant). .Human cut off: BH adjusted < 0.05, Mouse cut off: top 25 downregulated.

**Extended Data Figure 11:**
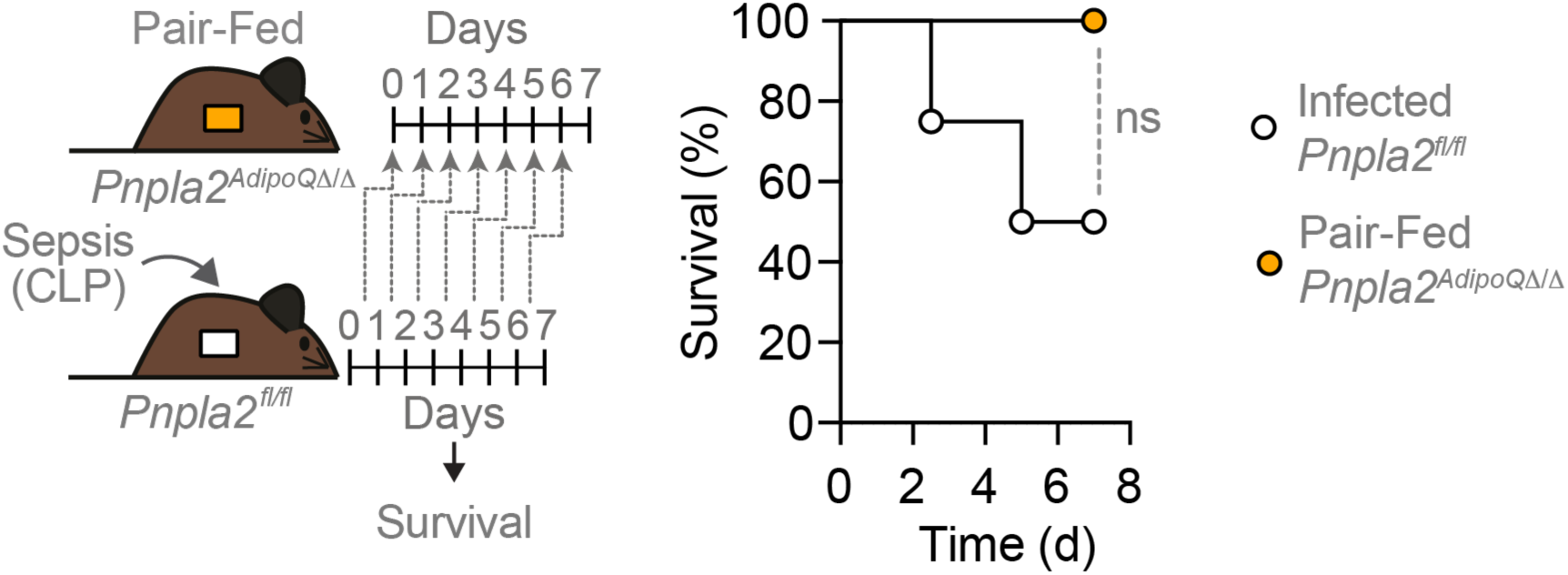
Illness induced anorexia is not sufficient to explain the lethal outcome of infection in *Pnpla2^AdipoQ11/11^* mice (Related to Figure 2). Survival of *Pnpla2^fl/fl^* mice (N=4) subjected to CLP (Sepsis), compared to pair-feed, uninfected *Pnpla2^AdipoQ11/11^* (N=4) mice. Log-rank (Mantel-Cox) test, ns- not significant.

**Extended Data Figure 12:**
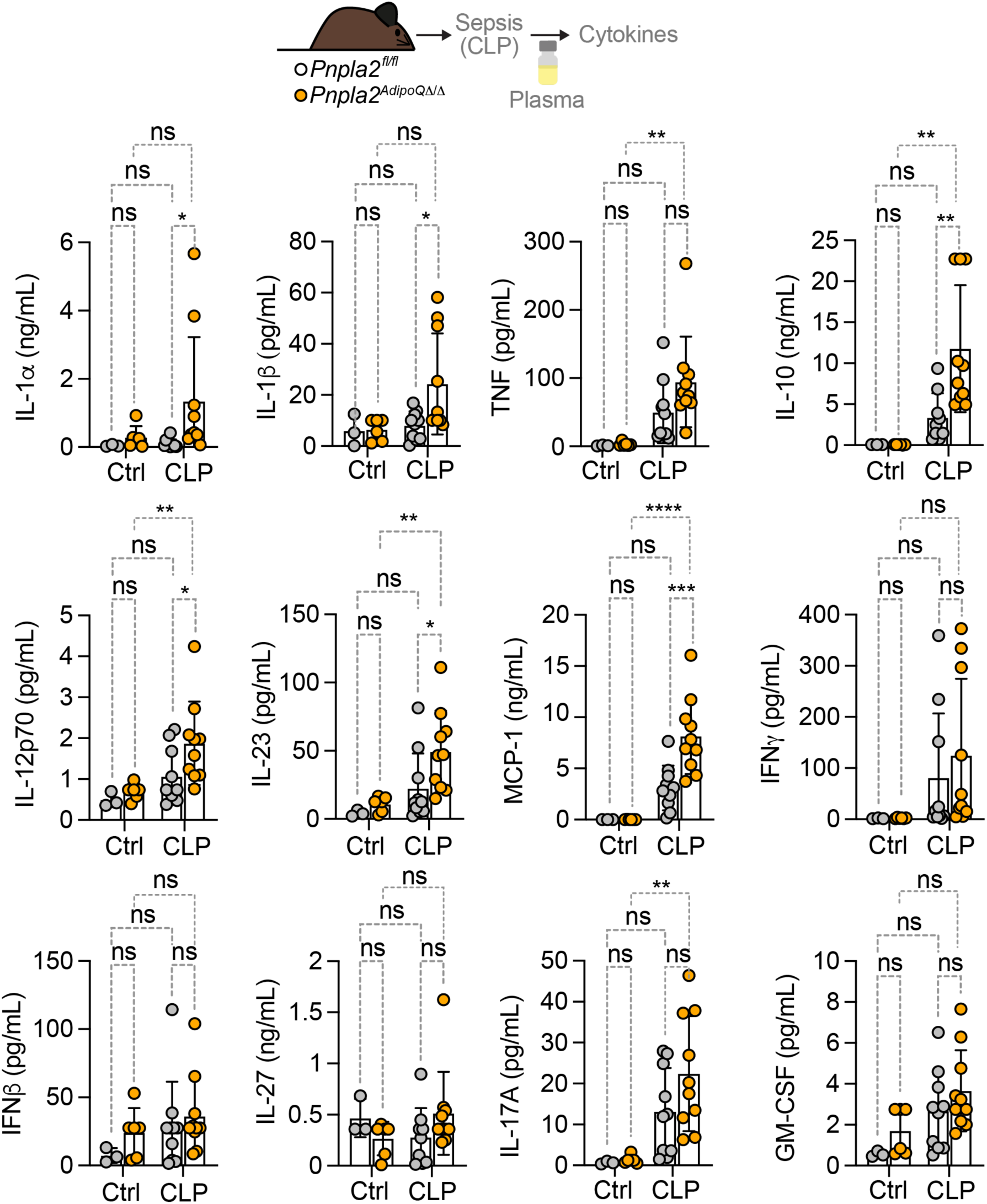
Adipocyte lipolysis modulates the cytokine response to infection in mice (Related to Figure 2). Cytokine concentration in plasma from naïve (control) *Pnpla2^fl/fl^*mice (N=3), naïve (Control, Ctrl) *Pnpla2^AdipoQ11/11^* (N=6) mice *vs.* septic *Pnpla2^fl/fl^* (N=10) and *Pnpla2^AdipoQ11/11^* (N=10) mice, 12hrs after CLP (Sepsis). Data represented as mean ± SD, pooled from 2 independent experiments, with similar trend. P-values calculated by Mixed-Effect-Model analysis with Šídák’s multiple comparisons test, ns- not significant, *p≤0.05, **p≤0.01, ***p≤0.001, ****p≤0.0001. Circles represent individual mice.

**Extended Data Figure 13:**
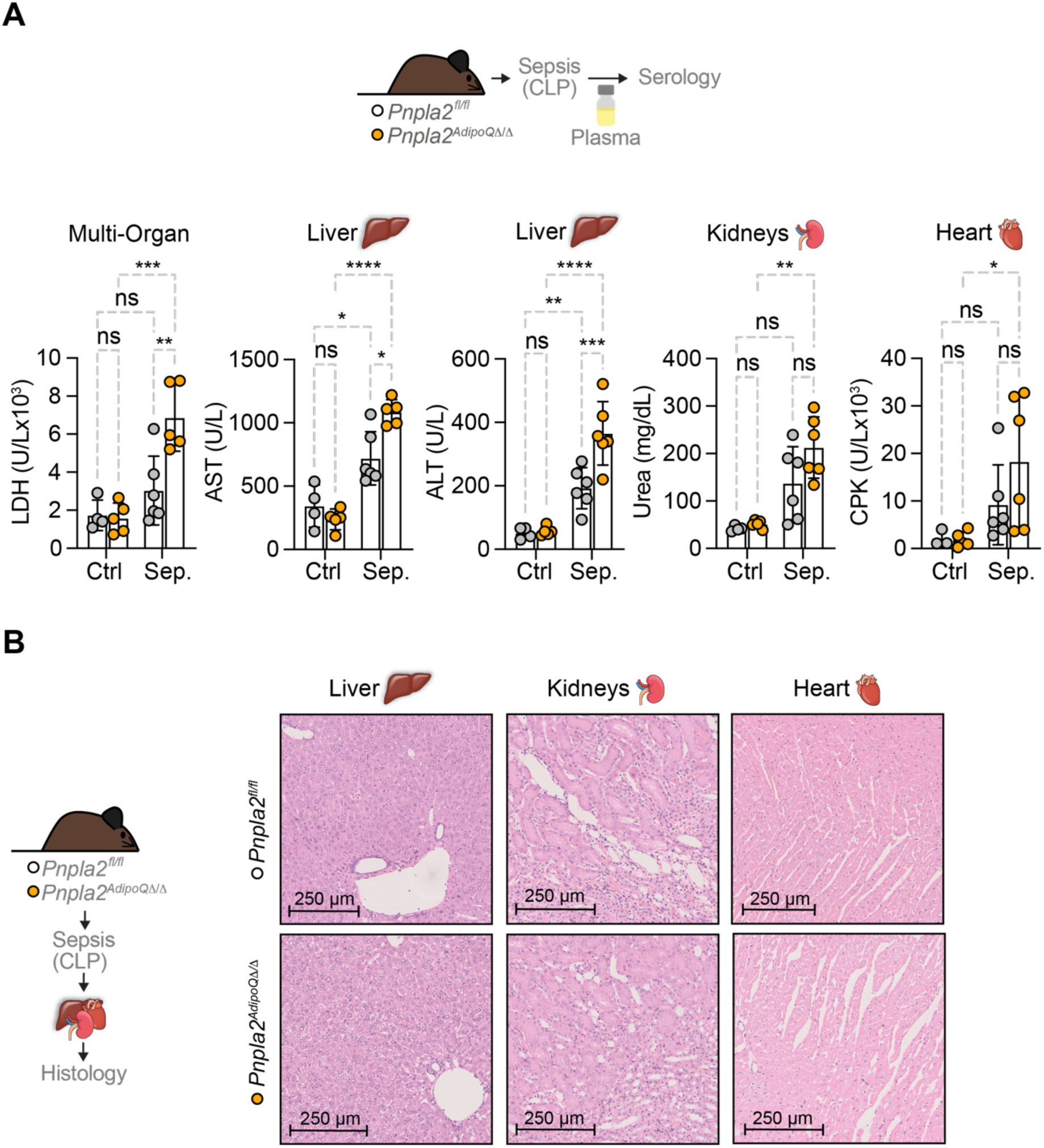
Adipocyte lipolysis protects infected mice from developing multi-organ damage (Related to Figure 2). **A)** Serology of organ damage markers: Lactate dehydrogenase (LDH; general tissue damage), Aspartate aminotransferase (AST; liver damage) and Alanine aminotransferase (ALT; liver damage), Urea (kidney damage) and Creatine phosphokinase (CPK; muscle/cardiac damage) from naïve (control) *Pnpla2^fl/fl^* mice (N=4), naïve (control) *Pnpla2^AdipoQ11/11^* (N=5) mice *vs.* septic *Pnpla2^fl/fl^* (N=6) and *Pnpla2^AdipoQ11/11^* (N=6) mice, 24hrs after CLP (Sepsis). Data represented as mean ± SD, pooled from 2 independent experiments, with similar trend. P-values calculated by Two-Way ANOVA with Tukeýs, ns- not significant, *p≤0.05, **p≤0.01, ***p≤0.001, ****p≤0.0001. Circles represent individual mice. **B)** Representative H&E staining of Liver, Kidney and Heart of septic *Pnpla2^fl/fl^* (N=3) and *Pnpla2^AdipoQ11/11^* (N=3) mice 24h after CLP (Sepsis).

**Extended Data Figure 14:**
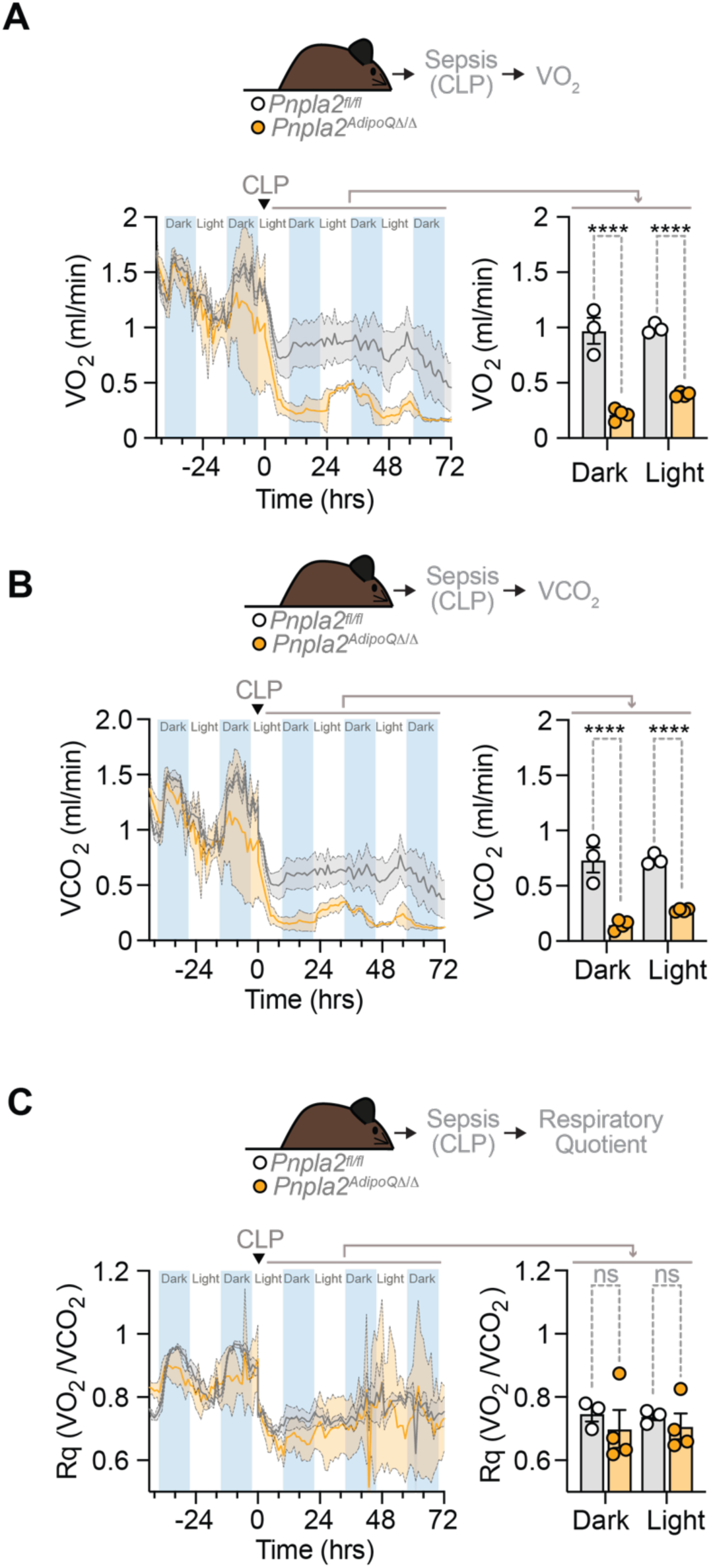
Adipocyte lipolysis supports energy metabolism in response to infection in mice (Related to Figure 2). **A)** Oxygen consumption (VO_2_) of *Pnpla2^fl/fl^* (N=4) or *Pnpla2^AdipoQ11/11^* (N=3) mice at steady state or subjected to CLP (Sepsis). Dotted line (0 h) indicates the induction of sepsis. Right panel shows the mean VO_2_ of daily averages at dark and light cycle, after CLP, as mean ± SEM. Circles represent individual mice. **B)** Carbon dioxide production (VCO_2_) in the same mice as A). Right panel shows the mean VCO_2_ of daily averages at dark and light cycle, after CLP, as mean ± SEM. Circles represent individual mice. **C)** Respiratory quotient (Rq) in the same mice as A). Right panel shows the mean Rq of daily averages at dark and light cycle, after CLP, as mean ± SEM. Circles represent individual mice. P-values in (A-C) calculated by Two-Way ANOVA with Šídák’s multiple comparisons test, ns- not significant, ****p≤0.0001.

**Extended Data Figure 15:**
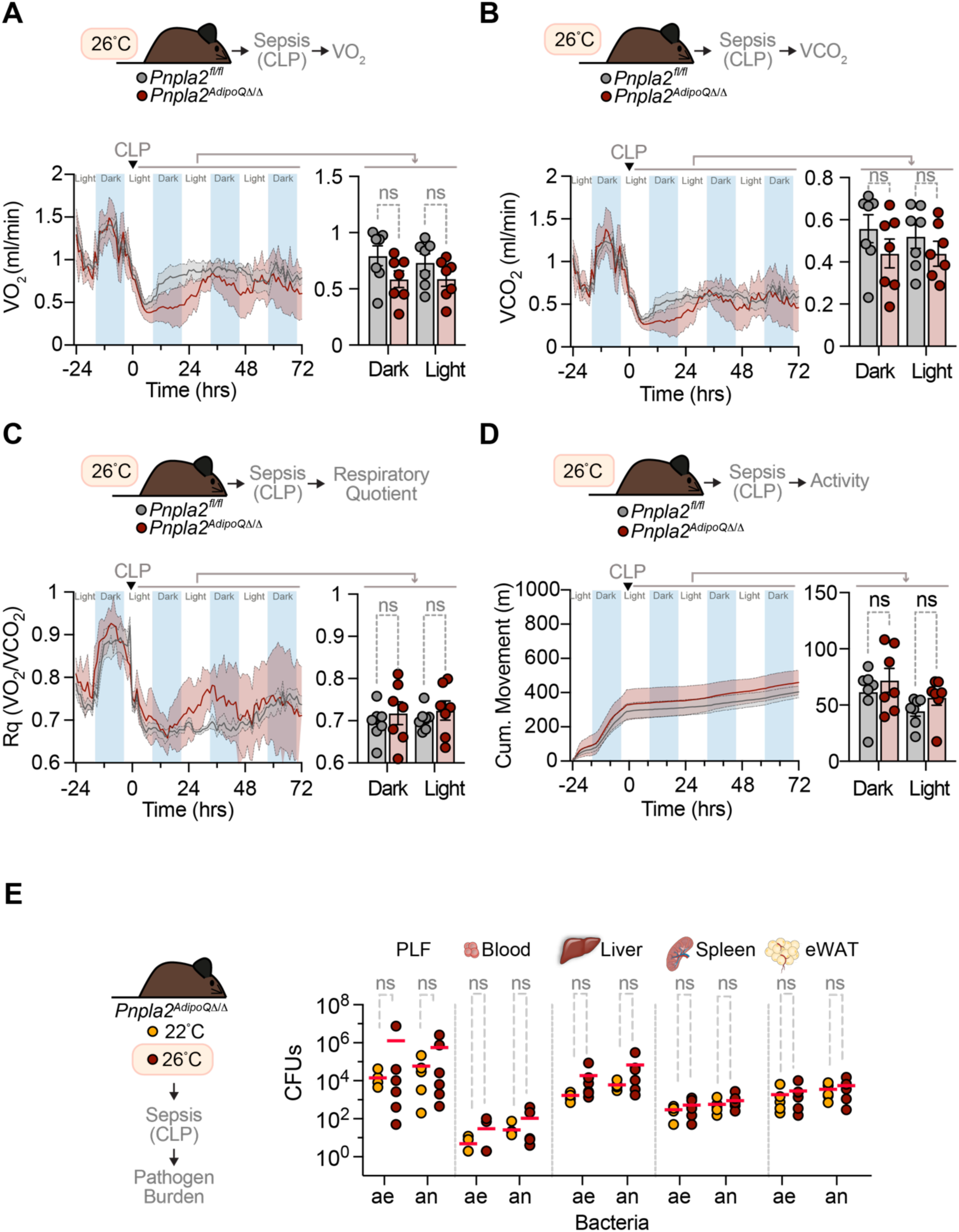
Adipocyte lipolysis supports an adaptive hypothermic response that establishes disease tolerance to infection in mice (Related to Figure 3). **A)**Oxygen consumption (VO_2_) of *Pnpla2^fl/fl^* (N=7) or *Pnpla2^AdipoQ11/11^* (N=7) mice at steady state or under septic (CLP) conditions. Ambient temperature of 26°C. Dotted line (0hrs) indicates the induction of sepsis. Right panel shows the mean VO_2_ of daily averages at dark and light cycle, after CLP, as mean ± SEM. **B)** Carbon dioxide production (VCO2) in the same mice as B). Right panel shows the mean VCO_2_ of daily averages at dark and light cycle, after CLP, as mean ± SEM. **C)** Respiratory quotient (Rq) in the same mice as B). Right panel shows the mean Rq of daily averages at dark and light cycle, after CLP, as mean ± SEM. **D)** Movement in the same mice as B). Right panel shows the mean movement of daily averages at dark and light cycle, after CLP, as mean ± SEM. B)-E) Data pooled from 2 independent experiments, circles represent individual mice. P-values calculated by Two-Way ANOVA with Šídák’s multiple comparisons test, ns- not significant. **E)** Colony forming units (CFU; ae: aerobe, an: anaerobe) from peritoneal lavage fluid (PLF), blood, liver, spleen and ependymal white adipose tissue (eWAT) of septic *Pnpla2^AdipoQ11/11^* mice at 22°C (N=5) or 26°C (N=6) ambient temperature (CLP; Sepsis, 24hrs). Data represented as mean (red bar), pooled from 2 independent experiments, with similar trend. Circles represent individual mice. P-values calculated by Students t-test on log-transformed values, ns-not significant.

**Extended Data Figure 16:**
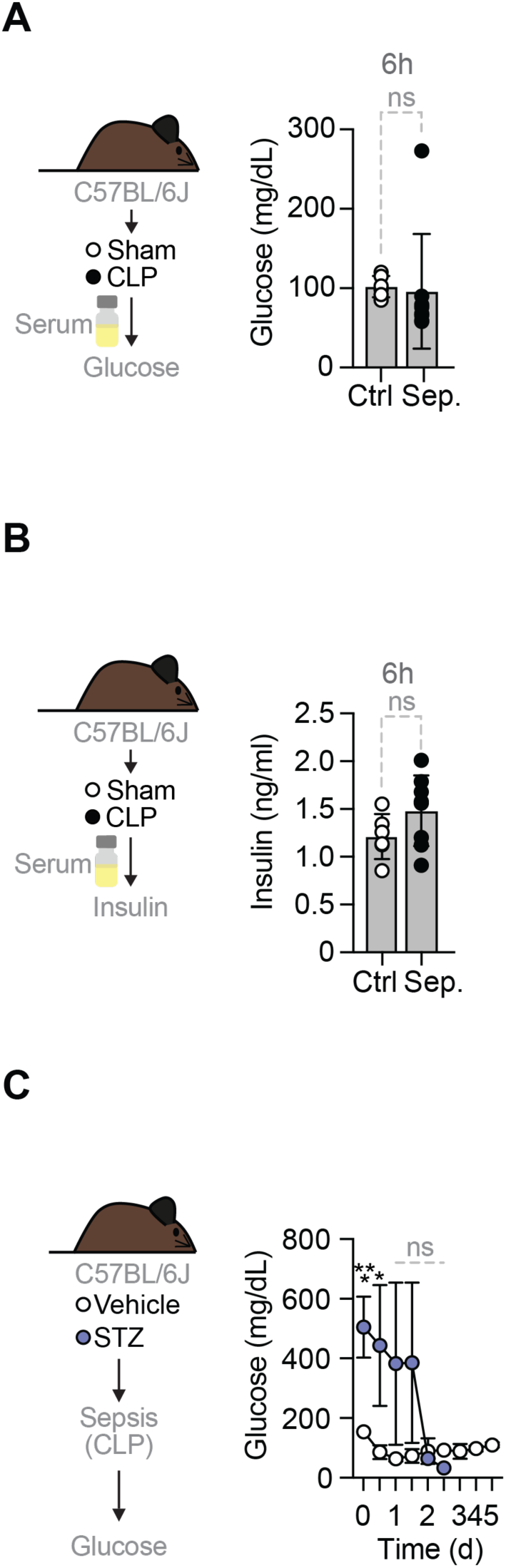
Glycemia and insulinemia in response to infection in mice. (Related to Figure 4). **A)** Blood glucose concentration of C57BL/6J mice, 6hrs after CLP (Sepsis; Sep.; N=8) or sham operation (Control; Ctrl. N=6). Circles represent individual mice. Plasma insulin concentration from C57BL/6J mice, 6hrs after CLP (Sepsis; Sep.; N=8) or sham operation (Control; Ctrl. N=6). A)-B) Data represented as mean ± SD, pooled from one experiment. P-values calculated by students t-test, ns-not significant. Circles represent individual mice. **C)** Blood glucose concentration of C57BL/6J mice, receiving PBS (vehicle; N=8) or Streptozocin (STZ; N=8) and subjected to CLP (Sepsis). Data represented as mean ± SD, pooled from 2 independent experiments. P-values calculated by mixed-effect analysis (Two-Way ANOVA) with Šídák’s multiple comparisons test, * p≤0.05; ***p≤0.001.

**Extended Data Figure 17:**
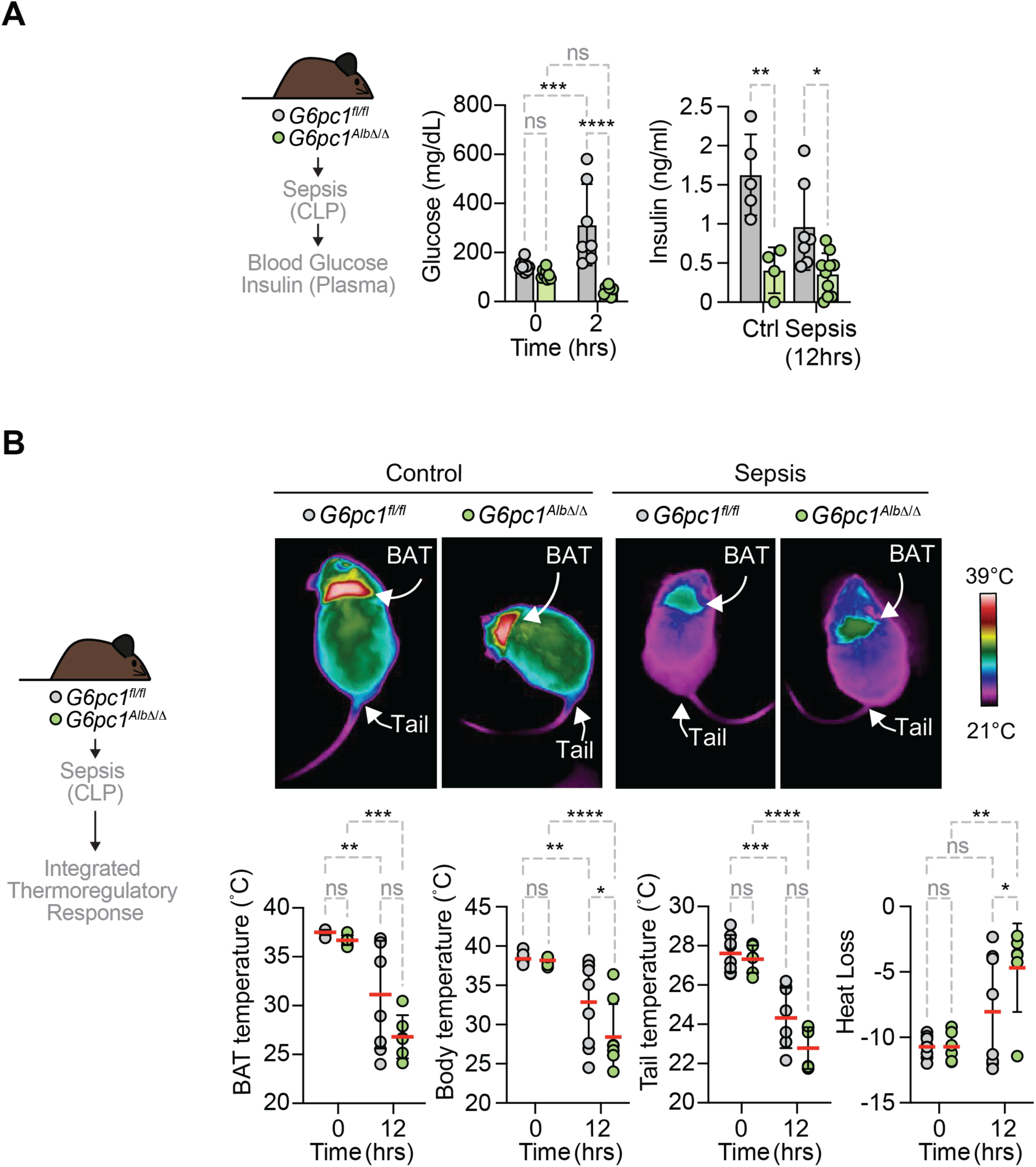
Hepatic glucose production regulates thermoregulation in response to infection in mice (Related to Figure 4). **A)** Blood glucose and plasma insulin concentration from *G6pc1^fl/fl^* or *G6pc1^Alb11/11^* mice subjected to CLP (Sepsis), at indicated time points, *vs.* control (Ctrl, *G6pc1^fl/fl^*: N=5; *G6pc1^Alb11/11^*: N=6-7). Data represented as mean ± SD, pooled from 2 independent experiments. P-values calculated by One-Way ANOVA with Tukeýs multiple comparison test, ns- not significant, *p≤0.05, **p≤0.01. **B)** Representative infrared images of *G6pc1^fl/fl^* or *G6pc1^Alb11/11^* mice subjected to CLP (Sepsis, 12hrs) or steady state (Controls, 0hrs). BAT, core body and tail temperatures as well as heat loss (ι1 of tail base and core body temperature). Data represented as mean ± SEM, averages from 23 different pictures of the same mouse (N=6-7 mice *per* genotype), pooled from 2 independent experiments, with similar trend. P-values calculated by Two-Way ANOVA with Šídák’s multiple comparisons test, ns- not significant, *p≤0.05; **p≤0.01; ***p≤0.001; ****p≤0.0001. Circles represent individual mice.

**Extended Data Figure 18:**
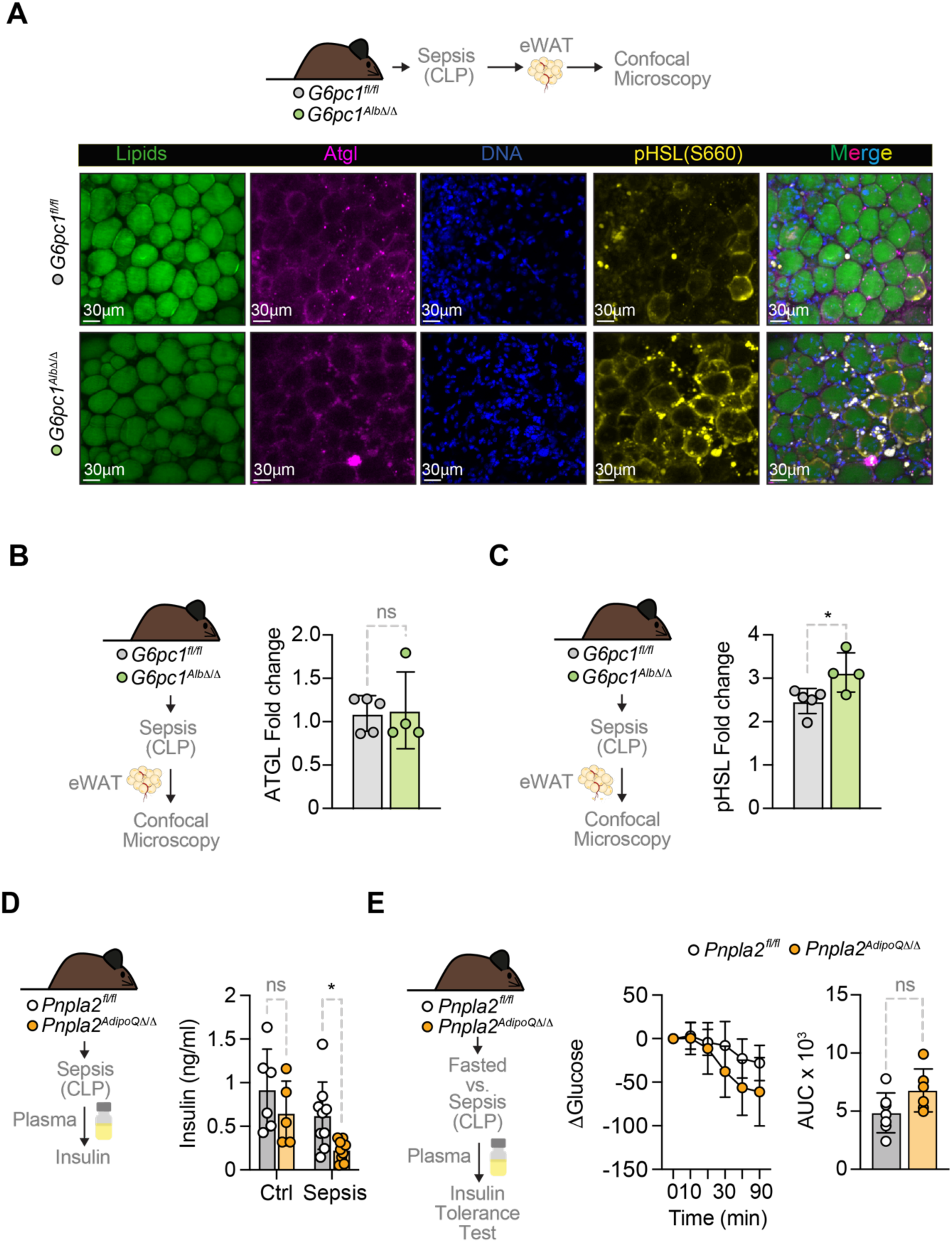
Adipocytes lipolysis sustains insulinemia in response to infection in mice (Related to Figure 4). **A)** Immunofluorescence imaging of neutral lipids (BodiPy; green), DNA (DAPI; blue), ATGL (anti-ATGL antibody with Alexa Fluor® 647; magenta) and phospho-HSL (anti-pHSL antibody with Alexa Fluor® 568; yellow) from the eWAT of *G6pc1^fl/fl^* and *G6pc1^Alb11/11^* 24hrs after CLP (Sepsis). **B)** Quantification of ATGL and **C)** p-Ser660-HSL (N=4-5 per group). Data represented as mean ± SEM, from 23 different areas in the same images, pooled from 23 images of 2 independent experiments with similar trend. Circles represent individual mice. P-values calculated by Student’s t-test, *p≤0.05. **D)** Plasma insulin concentration from *Pnpla2^fl/fl^*or *Pnpla2^AdipoQ11/11^* 12hrs after CLP (Sepsis) vs. control (N=6-9). Data represented as mean ± SD, pooled from 2 independent experiments, with similar trend. P-values calculated by Two-Way ANOVA with Tukeýs multiple comparison test, ns- not significant, *p≤0.05. **E)** Glycemia (left) and area under the curve (AUC, right) of *Pnpla2^fl/fl^*or *Pnpla2^AdipoQ11/11^* mice (N=6-7) injected with insulin 12hrs after CLP (Sepsis). Data represented as mean ± SD, 2 independent experiments. AUC analyzed by student’s t-test, ns - not significant. Circles represent individual mice.

